# Multimodal control of Cas13 activity through domain insertion at an allosteric hotspot

**DOI:** 10.1101/2025.04.21.649818

**Authors:** Liyuan Zhu, Long T. Nguyen, Alexandra G. Bell, Kara M. Gillmann, Harrison Oatman, Jack Hariri, Cameron Myhrvold, Jared E. Toettcher

## Abstract

CRISPR-Cas13 RNA nucleases have emerged as powerful tools for programmable RNA targeting. A light-controlled RNA nuclease could be transformative by enabling researchers to selectively knock down transcripts at desired positions in a cell or tissue or at timepoints of interest. Here, we develop a set of multimodal RfxCas13d tools that can be controlled by either light or small molecule addition. Screening an RfxCas13d library containing insertions of the AsLOV2 photoswitchable domain revealed an OptoCas13d-off variant that induced target RNA cleavage in the dark and switched to an inactive state under blue light. Insertion at this same allosteric hotspot could be further exploited to generate an OptoCas13d-on with the opposite light dependence and a ChemoCas13d that is activated upon the addition of rapamycin analogs. Through biochemical assays, we showed that AsLOV2 domain switching did not substantially affect Cas13d-RNA complex formation, indicating allosteric control over Cas13d catalytic activity. We applied the OptoCas13d-on system to target several endogenous transcripts and showed that it exhibited efficient mRNA knockdown only upon blue light illumination. Overall, our results demonstrate that engineered OptoCas13d can achieve cellular RNA modulation with high spatial and temporal precision.

## Introduction

Cas13, an RNA-targeting CRISPR effector protein, provides a versatile platform for programmable RNA cleavage, detection, editing and imaging^1–5^. RfxCas13d has been identified as one of the most efficient Cas13 orthologs for transcript knockdown in diverse cellular and animal contexts^1–4^. RfxCas13d binds to a cognate CRISPR RNA (crRNA). Pairing of the RfxCas13d-crRNA complex and a target RNA activates Cas13 nuclease activity, resulting in cleavage of the target RNA. Variations of this approach include using a catalytically dead Cas13 mutant fused to an RNA editing enzyme (e.g. ADAR2) to drive target RNA modification instead of degradation^5^.

One limitation with many current Cas13 tools is that they are constitutively active, performing RNA degradation or modification whenever Cas13, the crRNA, and the target RNA are present. More precise control over Cas13 activity would offer several advantages. First, RNA expression is precisely controlled in space and time to regulate functions both within cells (e.g., RNA trafficking in neurons) and across a tissue (e.g., gene expression in a developing embryo)^6–9^. Therefore, the development of a spatiotemporally controllable Cas13 system would enable more precise investigations into RNA function and regulation with high resolution. Additionally, recent studies have highlighted that Cas13 exhibits *trans*-cleavage activity: cleavage of RNAs other than the transcript of interest^10,11^. More precise control over the concentration of active Cas13 species could mitigate the collateral effects of *trans* cleavage and reduce cytotoxicity^12^.

Recent work has led to the development of split Cas13 systems including the padCas13^13^, CRISTAL (Control of RNA with Inducible SpliT Cas13 Orthologs and Exogenous Ligands)^14^, and an abscisic acid (ABA)-inducible split dCas13b system^15^. These approaches all rely on dual-component systems that function via the reconstitution of split proteins. Split proteins are powerful but can have several drawbacks, including the requirement to express two protein components that may each have reduced stability compared to the full-length protein. Split enzymes can also exhibit residual binding and leaky activity, as well as reduced catalytic activity once complemented^16,17^, which could limit the overall efficacy of target cleavage. Perhaps due to these drawbacks, the only light-controlled Cas13 tool developed to date works well as a nuclease-dead fusion to the ADAR base editor, but a catalytically active variant only achieves weakly light-switchable RNA knockdown^13^.

Here, we developed several single-domain controllable Cas13 nucleases based on domain insertion and allosteric control. Screening a library of AsLOV2 insertions into RfxCas13d revealed a single insertion site that conferred photoswitchable degradation of reporter RNA. Through subsequent optimization and insertion of distinct regulatory domains, we generated a light-inhibited variant via AsLOV2^18,19^ insertion (OptoCas13d-off), a light-activated variant through LightR insertion^20^ (OptoCas13d-on), and a rapamycin-inducible variant incorporating the UniRapR^21^ domain (ChemoCas13d). We further validated the efficacy of light-activated OptoCas13d-off in mediating light-inducible knockdown of endogenous transcripts, highlighting its potential as a powerful tool for studying gene function with high spatiotemporal resolution. These findings underscore the effectiveness of combinatorial screening approaches in identifying allosteric hotspots within structurally complex proteins, paving the way for future advances in optogenetic, chemically inducible, and other modalities of allosterically controlled protein engineering.

## Results

### Construction of RfxCas13d-AsLOV2 random insertion library and reporter landing pad cell line

In recent years, insertion of an optogenetic or chemogenetic domain into a protein of interest has gained prominence as a technique for creating stimulus-switchable proteins^19,20,22,23^. However, finding suitable insertion sites can be difficult, particularly for large proteins where structural information is incomplete. We recently reported that domain insertion profiling^24^ and screening could be used for unbiased discovery of suitable insertion sites^25^. We generated a library in which the AsLOV2 photoswitchable domain was inserted at many random positions into a Gal4-VP64 transcription factor and screened variants for photoswitchable gene expression, yielding an OptoGal4 transcription factor with a >150-fold change in gene expression between dark and illuminated states^25^. However, domain insertion has not yet been attempted for large, complex enzymes such as RfxCas13d (∼1,000 amino acids), which possess multiple functional domains and are structurally less well-characterized.

We first set out to construct a comprehensive library of AsLOV2 insertions into the RfxCas13d protein scaffold, and to establish a cell line in which to test the efficacy of this library for RNA knockdown in mammalian cells (**Figure 1a, b**). For library construction, we employed a well-adopted approach for generating a random domain insertion library, referred to as DIP-Seq^24^ (**Fig. 1c**). This approach utilizes a Mu transposase to randomly insert the chloramphenicol resistance (CmR) gene into a target gene of interest. The chloramphenicol-resistant library is then inserted into an expression vector, followed by replacement of CmR with the domain of interest. We modified the original DIP-Seq protocol to adjust for the insertion of a small domain into large target proteins and to account for the presence of BsaI sites in the destination vector (**Supplementary Fig. 1**). After moving the library of AsLOV2-inserted RfxCas13d variants into the final landing pad recombination vector, we performed next-generation DNA sequencing of the library and showed that our library covers at least 91% of possible insertion sites (**Fig. 1d**). The resulting library was then integrated into a reporter cell line for follow-up screening.

**Figure 1.**
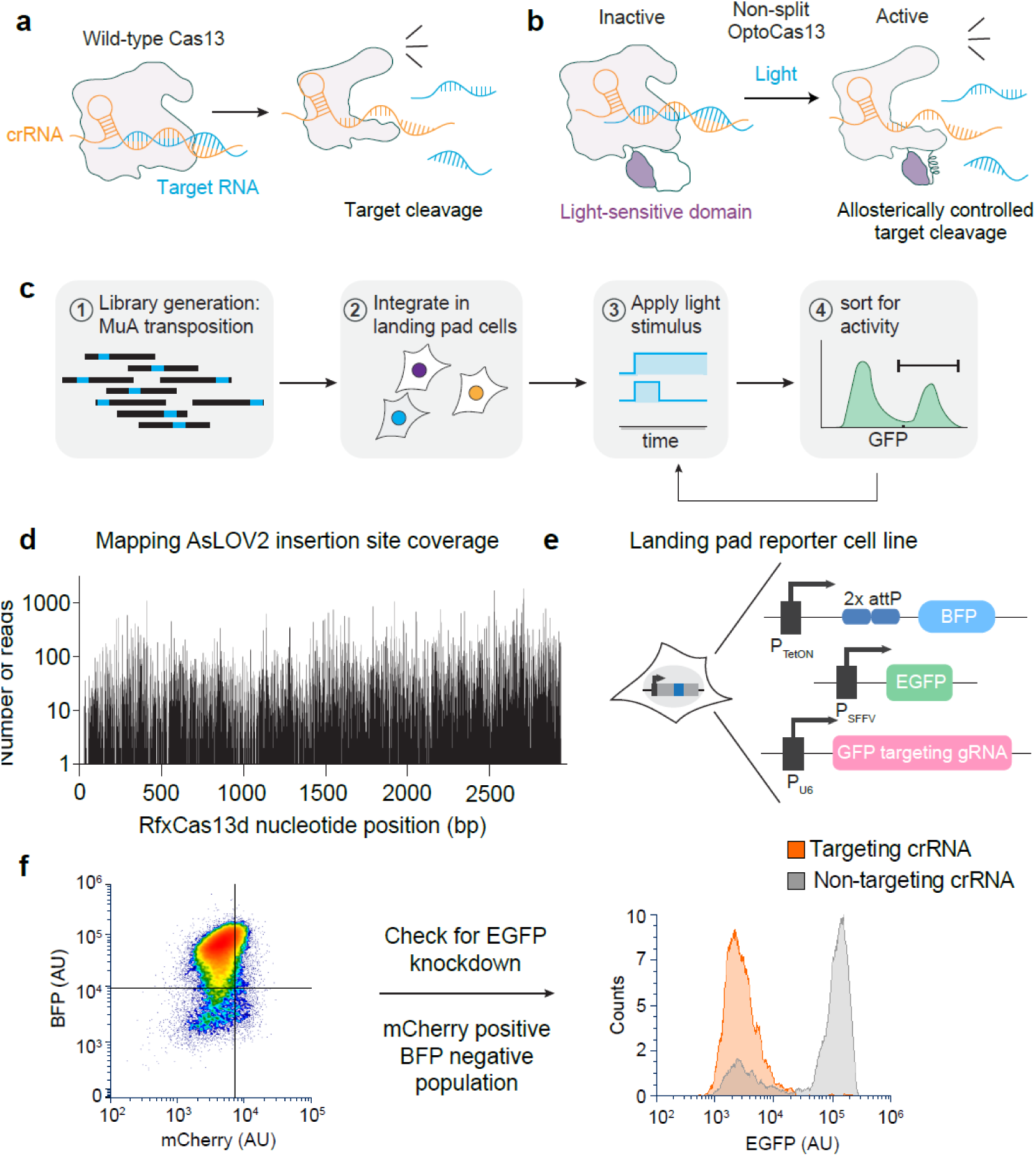
Engineering approach for generating optogenetically controllable RfxCas13d variants. (**a**) and (**b**) Comparison of mechanism of a wild-type RfxCas13d cleavage and an allosterically controlled RfxCas13d via light-sensitive domain insertion. (**c)** Schematic of the workflow for generating RfxCas13d-AsLOV2 insertion library and incorporating them into landing pad reporter cell lines for selections. **(d)** Coverage of random AsLOV2 insertion sites in between amino acid residues within RfxCas13d through next-generation sequencing analysis. **(e)** Composition of a landing pad reporter cell line. This cell line contains a single recombination site with BFP gene in frame for negative selection. This recombination site acts as a landing pad for the incorporation of the RfxCas13d-AsLOV2 insertion library. The landing pad cell line also contains a reporter EGFP gene and a crRNA targeting EGFP mRNA. (**f**) Successful incorporation of the RfxCas13d-AsLOV2 insertion library results in displacement of the BFP gene by mCherry. BFP negative and mCherry positive cell population was sorted via fluorescence-activated cell sorting (FACS). EGFP knockdown was confirmed in this sorted subpopulation.

We next created a HEK293T landing pad reporter cell line where the activity of RfxCas13d could be assessed by EGFP knockdown, allowing for the measurement and screening of RfxCas13d library variants through fluorescence-activated cell sorting (FACS)^26,27^ (**Fig. 1e**). The reporter cell line was constructed to include both a constitutively expressed EGFP reporter and an EGFP mRNA-targeting CRISPR RNA (crGFP), so that the introduction of a functional RfxCas13d variant could cleave EGFP mRNA and reduce the cell’s overall EGFP expression. This landing pad cell line allows for the generation of a pooled library of RfxCas13d insertion variants, with each cell expressing a single variant from a defined genomic locus in the cell (**Fig. 1e**). By sorting cells that are mCherry-positive and BFP-negative after transfection, we could enrich only those cells with successful landing pad incorporation. A ∼100-fold decrease in EGFP expression was observed in RfxCas13d-expressing reporter cells compared to an analogous cell line expressing a non-targeting crRNA, consistent with robust RfxCas13d/crRNA-targeted EGFP degradation in this cell line (**Fig. 1f**).

### Domain insertion at the QK634 allosteric hotspot confers light-switchable Cas13 activity

We introduced the RfxCas13d-AsLOV2 library into the landing pad cell line and set out to screening the library for EGFP knockdown in both light and dark conditions (**Fig. 2a**). We observed that after the library was initially expressed, many variants exhibited high EGFP levels with a small subpopulation still successfully inducing EGFP knockdown. However, a major challenge arose—these knockdown-competent populations were lost after approximately three weeks in culture, indicating a potential selection against continuous Cas13 activity in the landing pad cell line (**Fig. 2b**). We hypothesized that constitutive expression of RfxCas13d may have been toxic in this cell line due to the high expression level of RfxCas13d and/or *EGFP* target RNA driving *trans* cleavage activity^12^.

**Figure 2.**
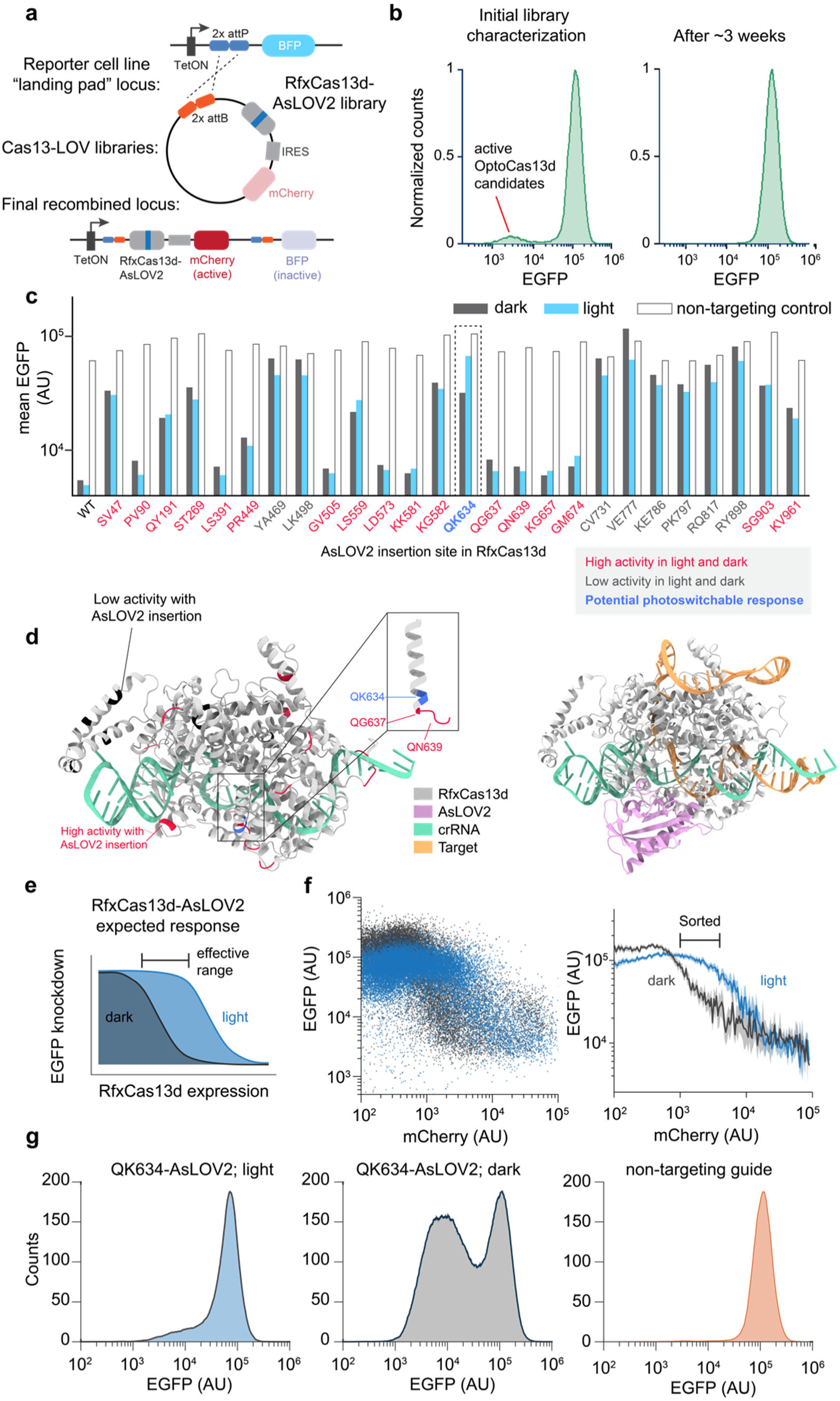
Screening RfxCas13d-AsLOV2 variants for light switchable behavior. (**a**) Detailed schematic of successful recombination of RfxCas13d-AsLOV2 insertion library into the landing pad reporter cell line. (**b**) Histogram showing the loss of EGFP knockdown during routine cell culture, indicating a potential selection against active variants. (**c**) Individual candidates were tested for photo-switchability. Many variants either exhibited loss of function due to domain insertion or no switchable behavior. Site QK634 was identified as a potential candidate. (**d**) Predicted structure of wild-type RfxCas13d binary and RfxCas13d-QK634-AsLOV2 tertiary complex with 27 sites highlighted. crRNA contains a spacer targeting EGFP mRNA. Prediction was performed using AlphaFold 3. **(e)** Schematic of effective range of RfxCas13d-QK634-AsLOV2 variant photo-switchability. **(f)** Flow cytometry indicating single-cell expression of EGFP (indicating RfxCas13d functionality) and mCherry (indicating RfxCas13d expression level) under light and dark conditions. The effective range for a light-controlled response is the range of mCherry levels showing distinct EGFP expression in light versus dark. Right panel: error bars shown mean + 95% confidence interval for single cells binned from **f**. Data shown is from one experiment representative of n=3 replicates. **(g)** Gated cells were tested for knockdown of EGFP upon transfection with EGFP-targeting guide RNA.

Faced with this setback, we hypothesized that the set of highly active variants that were lost during culture might also include variants with photoswitchable activity, since these variants tolerated AsLOV2 insertion while remaining active in the dark. To identify the insertion sites represented by these variants, we performed next-generation sequencing of the library before and after cell incorporation. While most insertion sites were present at similar proportions in both libraries, we identified 27 sites that exhibited clear depletion after incorporation into cells and continuous culture (**Supplementary Fig. S2c**). We selected these variants for individual testing to assess their knockdown activity under dark or light conditions. Among the 26 OptoCas13d variants tested (one variant was not recovered during cloning), 16 retained constitutive high activity under both dark and light conditions, and 9 showed low activity in both dark and light. Notably, we identified one OptoCas13d variant with an insertion at site QK634 (AsLOV2 inserted between Glu634 and Lys635) which exhibited stronger EGFP knockdown in the dark compared to the light (**Fig. 2c** and **Supplementary Fig. 3**).

To better understand the structural context of each insertion site, we modeled both binary (RfxCas13d:crRNA) and tertiary (RfxCas13d:crRNA:target RNA) complexes using AlphaFold 3^28^ (**Fig. 2d and Supplementary Fig. 4**). The AlphaFold 3 predictions indicated that some insertions were positioned within well-folded structural motifs (e.g. within α-helices or β-strands of the protein), providing a likely explanation why AsLOV2 insertion in these positions could break Cas13d functionality. We then mapped the potential allosteric hotspot, QK634 (highlighted in green), onto the predicted structure of RfxCas13d. This site is situated within the Helical-2 linker domain, which bridges two catalytic domains. Structural predictions indicate that QK634 is surface-exposed and positioned spatially distant from the active sites. A closer examination of the QK634 position revealed that it resides at the transition point between an unstructured loop— where AsLOV2 insertion does not disrupt Cas13 functionality in either dark or light conditions (see e.g., the QG637 and QN639 sites)—and a well-folded α-helix, where AsLOV2 insertion completely abolishes enzymatic activity. This unique positioning may enable QK634 to accommodate AsLOV2 insertion while effectively transmitting the conformational changes induced by light to regulate overall OptoCas13d functionality (**Fig. 2d and Supplementary Fig. 4**).

We next sought a more quantitative picture of light-switchable EGFP knockdown in the QK634 RfxCas13d variant, henceforth termed OptoCas13d-off. We reasoned that the conformational switch might exhibit concentration dependence, switching the threshold of Cas13d expression at which potent EGFP knockdown would be observed between light and dark cases (**Fig. 2e**). We analyzed EGFP reporter expression in cells transfected with an OptoCas13d-off IRES mCherry construct, so that the OptoCas13d-off expression level could be indirectly visualized using mCherry expression. Indeed, we observed that EGFP knockdown in light and dark depended on the Cas13 expression level. This experiment also led us to identify a specific mCherry expression range where EGFP knockdown occurred in the dark but not in cells illuminated with blue light (**Fig. 2f**).

We next generated a clonal EGFP-expressing cell line harboring the OptoCas13d-off IRES mCherry and sorted for the specific mCherry expression level where we observed light-inducible knockdown (**Supplementary Fig. 6a**). Unlike the landing pad reporter cell line (**Fig. 2a**), which stably integrates both EGFP and an EGFP-targeting crRNA, this clonal cell line did not include a crRNA, ensuring that Cas13 incorporation would not affect cell viability. Upon transfection of this clonal cell line with an EGFP-targeting crRNA, we observed EGFP knockdown comparable to wild-type Cas13 activity in dark conditions, whereas knockdown was nearly absent under light exposure or upon transfection with a non-targeting crRNA (**Fig. 2g**). Collectively, these findings demonstrate that inserting AsLOV2 at the QK634 site in RfxCas13d yields a light-inhibited Cas13d variant.

### Biochemical and cellular assays reveal mechanistic basis for light-switchable RfxCas13d activity

We next set out to gain insight into the mechanism of light-gated RNA cleavage using biochemical assays. We reasoned that three different processes could in principle be regulated by the photoswitchable domain: RfxCas13d:crRNA binary complex formation, RfxCas13:crRNA:target RNA ternary complex formation, or RfxCas13d cleavage of the target RNA.

We expressed and purified active and catalytically inactivated mutants of wild-type RfxCas13d and OptoCas13d-off in *E. coli*. First, we performed *in vitro trans*-cleavage assays, as previously described^29^, to determine if light-dependent cleavage could also be observed *in vitro*. For consistency, we used the same EGFP crRNA and synthetic mRNA fragment to assess *in vitro* cleavage. *Trans*-cleavage activity for wild-type RfxCas13 and OptoCas13d-off were measured immediately after exposing the RNP complex to blue (∼465 nm) light or dark treatment followed by the addition of target RNA (**Fig. 3a**). Notably, we noticed a substantial reduction in relative fluorescence for OptoCas13d-off under blue light compared to its dark treatment counterpart, corroborating the same observation for EGFP knockdown in HEK293T cells. On the other hand, the wild-type RfxCas13d exhibited negligible differences in fluorescence intensity between dark and blue light conditions (**Fig. 3b,c**). These data confirm that OptoCas13-off retains photoswitchable cleavage activity *in vitro*.

**Figure 3.**
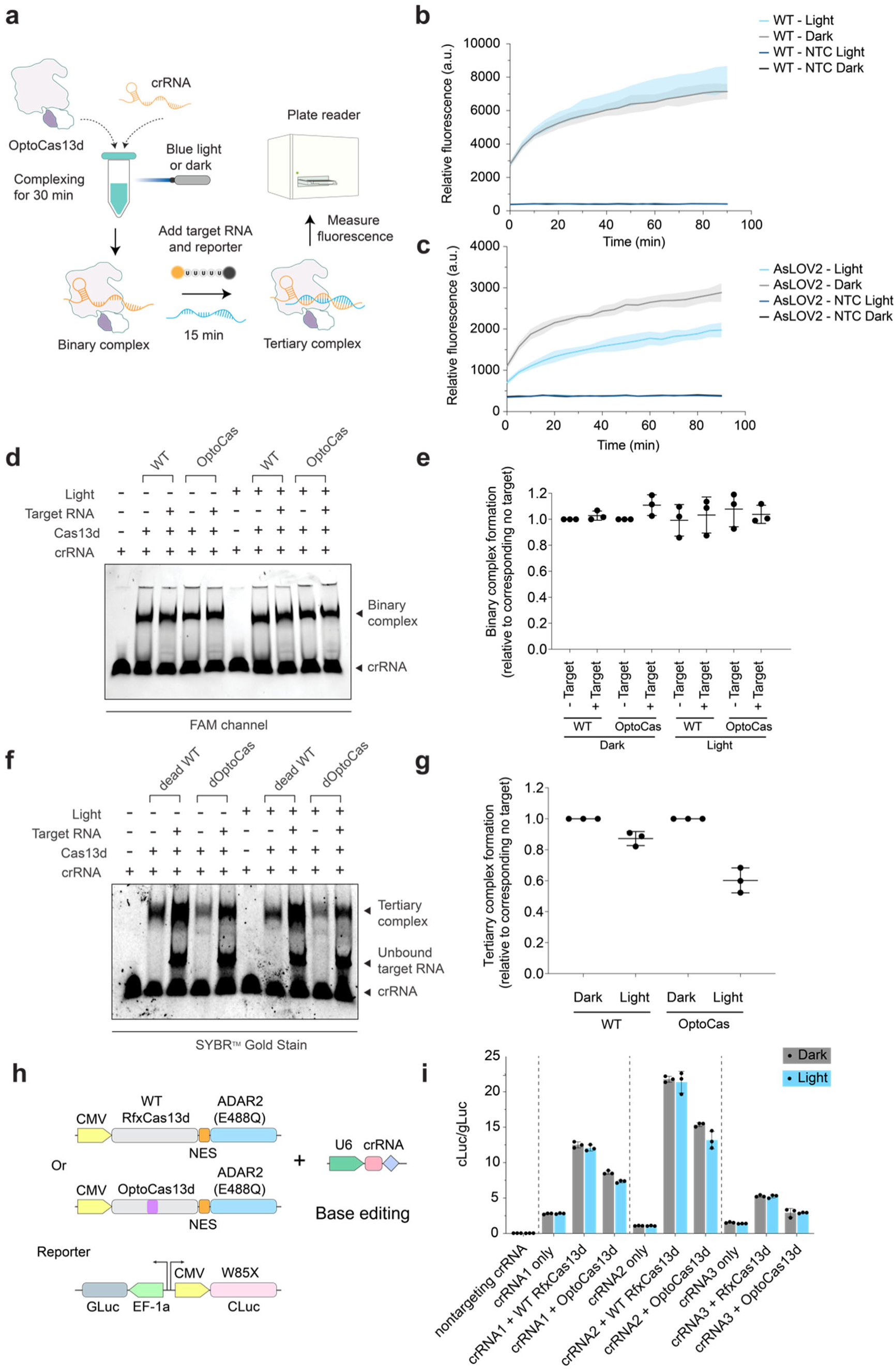
*In vitro* biochemical characterization of RfxCas13d-QK634-AsLOV2 variant. (**a**) Schematic of trans-cleavage assay workflow with the incorporation of a light treatment step. (**b**) and (**c**) Comparison of *in vitro* cleavage activity of wild-type RfxCas13d and RfxCas13d-QK634-AsLOV2 in dark and light conditions. Error bars in **b-c** show mean + SD for 3 independent replicates. (**d**) EMSA assay of binary complexation of wild-type RfxCas13d and RfxCas13d-QK634-AsLOV2 in dark and light conditions. (**e**) Quantification of band intensity in (**d**). (**f**) EMSA assay of tertiary complexation of wild-type RfxCas13d and RfxCas13d-QK634-AsLOV2 in dark and light conditions. (**g**) Quantification of band intensity in (**f**). (**h**) Schematic of wild-type RfxCas13d and RfxCas13d-QK634-AsLOV2 constructs used for luciferase base editing. (**i**) A-to-G conversion of W85X in the CLuc mRNA for wild-type RfxCas13d and RfxCas13d-QK634-AsLOV2 in light and dark conditions. Error bars show mean + SD from n=3 independent replicates.

To test whether light might regulate the formation of OptoCas13-off binary or tertiary RNA complexes, we turned to electrophoretic mobility shift assays (EMSAs) to measure changes in RNA-protein binding. We incubated OptoCas13-off with an EGFP-targeting crRNA whose 3’ end was labeled with a FAM fluorophore to measure changes in crRNA:Cas13 binary complex formation and found that blue light illumination had no effect on the fraction of crRNA:Cas13 binary complexes for either wild-type RfxCas13d or OptoCas13d-off (**Fig. 3d,e** and **Supplementary Fig. 5**). For analysis of tertiary complex formation, we purified dead Cas13 variants of (dRfxCas13d and dOptoCas13d-off) harboring R239A/H244A/R858A/H863A mutations that allow for RNA binding but not cleavage. We observed a modest decrease in the amount of target RNA binding for OptoCas13d-off under blue light illumination compared to the dark-incubated complex (**Fig. 3f,g**). Together, these results suggest that blue light does not affect the binary complex formation but at least partially alters the binding of crRNA:RfxCas13d to its target. However, the modest effect on tertiary complex formation suggested light might also affect OptoCas13d-off RNA nuclease activity.

We constructed an alternative Cas13-based system to directly distinguish light-dependent effects on RNA binding or catalysis in cells. It was recently shown that a dead Cas13 variant could be fused to the ADAR2 base editor to perform RNA-guided base editing^28^. We thus reasoned that an ADAR2-fused dead OptoCas13d-off would exhibit light-dependent base editing only if light primarily acted on Cas13/mRNA complex formation. Alternatively, if light stimulation altered catalytic activity, base editing would be unaffected by optogenetic stimulation. To distinguish between these models, we generated catalytically inactive versions of wild-type RfxCas13d (dRfxCas13d) and its light-controlled variant (dOptoCas13d-off) by fusing them to the ADAR2-based RNA base editor, as described by Cox *et al*^5^. These constructs were evaluated using a dual-luciferase reporter system, in which a non-functional CLuc was used as the target, and GLuc served as an internal normalization control. The intended A-to-G conversion was designed to correct a premature stop codon at amino acid position 85 of the CLuc gene (**Fig. 3h**)^5^. Consistent with our hypothesis, we observed minimal differences in base editing efficiency between dark and light conditions, with both exhibiting reduced editing activity compared to wild-type RfxCas13d (**Fig. 3i**). These findings aligned with the results obtained from biochemical assays, further supporting the notion that OptoCas13d-off relies primarily on light-gated RNA cleavage activity, providing a potent switch for controlling RNA degradation but not influencing dead-Cas13-based tools such as RNA base editors.

### Development of multimodal RfxCas13d variants with alternative regulatory domains

Although our OptoCas13d-off system confers potent light-dependent regulation of RNA cleavage, we can envision applications where alternative stimulus modalities would be preferable to induce Cas13 activity. Fortunately, the principle of allosteric control is quite modular, permitting multiple light- and drug-switchable domains to be in principle inserted at the same position. To enable multimodal control of RfxCas13d activity, we replaced the AsLOV2 domain at the QK634 site with alternative domains capable of undergoing conformational changes in response to specific stimuli.

To demonstrate the modularity of our approach, we selected two domains — UniRapR and LightR — both of which have been previously reported to confer stimulus-inducible properties to target proteins upon insertion^20,21^. UniRapR is a single-chain protein composed of a modified insertable FKBP subdomain fused to two helices derived from FRB. Upon rapamycin binding, UniRapR undergoes an intramolecular conformational switch by bringing its two subdomains into proximity. In contrast, LightR consists of two VVD domains linked by a flexible peptide linker^20^. VVD is a light-sensitive domain that exists as a monomer in the absence of light but undergoes homodimerization upon light exposure, leading to an overall conformational change in LightR. Structurally, AsLOV2 exists in a closed conformation in the dark, with its two terminal loops positioned proximally. Light exposure induces undocking of these loops, transitioning AsLOV2 into an open conformation^30^. In contrast, UniRapR adopts an open conformation under normal conditions but undergoes closure upon rapamycin addition, while LightR remains open in the dark and transitions to a closed conformation in response to light. Based on these properties, we hypothesized that substituting AsLOV2 with UniRapR or LightR in RfxCas13d would render the enzyme responsive to rapamycin or light, respectively, but with stimulus-dependent activation rather than inhibition as is the case for AsLOV2 (**Fig. 4a**).

**Figure 4.**
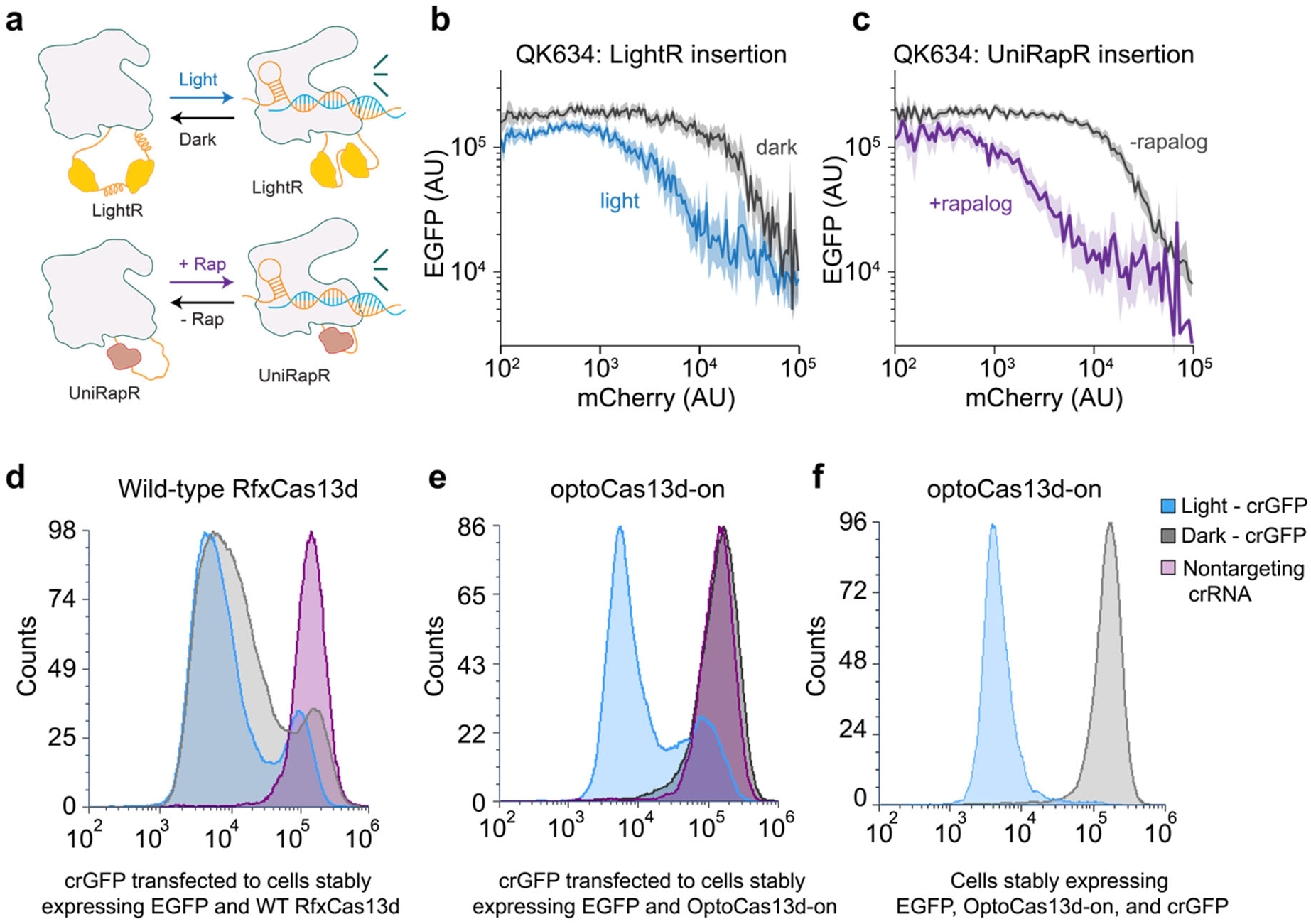
Expansion of controllable RfxCas13d variants with alternative regulatory domains. **(a)** Schematic depicting LightR and UniRapR domains inserted between residues QK634 in RfxCas13d. **(b)** RfxCas13d-QK634-LightR demonstrates light inducible knockdown EGFP. **(c)** RfxCas13d-QK634-UniRapR demonstrates chemical inducible knockdown of EGFP. For **b-c**: error bars shown mean + 95% confidence interval for single cells in flow cytometry experiment. Data shown is from one experiment representative of n=3 replicates. **(d)** and **(e)** Knockdown of EGFP where crRNA is transfected in and expression of effector proteins is under a doxycycline inducible promoter for wild type RfxCas13d and RfxCas13d-QK634-LightR, respectively. **(f)** Histogram showing robust EGFP knockdown (∼100-fold) of cells that stably express EGFP, RfxCas13d-LightR and crRNA targeting EGFP mRNA with light treatment.

To test this hypothesis, we transfected an EGFP-expressing stable cell line with RfxCas13d variants tagged with IRES-mCherry and co-transfected an EGFP mRNA-targeting crRNA. As anticipated, insertion of LightR into QK634 conferred light-induced activation of RfxCas13d (termed OptoCas13d-on; **Fig. 4b**), whereas UniRapR insertion resulted in rapamycin-dependent activation (termed ChemoCas13d; **Fig. 4c**). As in the case of OptoCas13d-off, both variants were only stimulus sensitive at intermediate expression levels, although this range was larger for OptoCas13d-on and ChemoCas13d compared to the original OptoCas13d-off, possibly due to a larger stimulus-induced conformational change of these variants.

We generated a clonal OptoCas13d-on cell line within the stimulus-switchable expression range to explore this system’s full potential for light-switchable RNA regulation. To mitigate any potential toxicity, we employed a doxycycline-inducible system for controlled expression. The RfxCas13d-LightR coding sequence, tagged with IRES-mCherry, was cloned downstream of a Tet-On promoter in a piggyBac vector. This construct was transfected into a 293T landing pad cell line expressing a destabilized EGFP-PEST (293T-LP-dGFP) along with a piggyBac helper plasmid to facilitate genomic integration. Notably, 293T-LP-dGFP cells already stably express the reverse tetracycline-controlled transactivator (rtTA) from the landing pad locus, enabling transcriptional activation of RfxCas13d-LightR in response to doxycycline induction.

We performed clonal selection following transfection and doxycycline treatment, focusing on cells exhibiting mCherry fluorescence within the effective expression range (∼10⁴ fluorescence units). To validate the functionality of this stable cell line, we transfected it with EGFP-targeting crRNA. As expected, EGFP knockdown was observed exclusively under blue light exposure, with no detectable knockdown in the dark (**Fig. 4d, e**), demonstrating that OptoCas13-on can act as a potent light-activated RNA expression switch. Stable integration of the EGFP-targeting crRNA via lentiviral transduction produced an even more homogeneous EGFP knockdown response to light (**Fig. 4f**). Overall, these experiments demonstrate that the OptoCas13 and ChemoCas13 tools are single-protein systems with potent, stimulus-switchable, and programmable RNA cleavage.

### Applications of OptoCas13d-on for light-inducible endogenous transcript knock-down

Thus far we have successfully developed OptoCas13 and ChemoCas13 systems for stimulus-controlled knockdown of an EGFP reporter. However, the ultimate test of such a tool is whether it can be used for stimulus-controlled knockdown of endogenous RNAs. We focused on OptoCas13d-on, as light-inducible gene silencing is a desirable functionality for many applications and this system offers significant potential for studying gene function with high temporal and spatial resolution. We transfected the doxycycline-inducible OptoCas13d-on cell line with a guide RNA (crRNA) targeting the endogenous transcript B4GALNT1. Cells were either exposed to light for 5 hours following transfection or maintained in darkness. After an additional 32 hours, total RNA was extracted, and RT-qPCR was performed to quantify B4GALNT1 mRNA levels (**Fig. 5a**). As expected, a significant knockdown of B4GALNT1 was observed exclusively in light-exposed samples, confirming the light-dependent activity of OptoCas13d-on. Given that the presence of non-transfected cells might reduce the observed knockdown efficiency, we anticipated that generating a clonal cell line stably expressing both OptoCas13d-on and the crRNA could enhance the effect still further (see e.g., **Fig. 4i**). We further evaluated the performance of OptoCas13d-on in silencing five additional endogenous transcripts by transfecting our cell line with multiple crRNAs targeting AXNA4, FTH1, CD99, CLTA, and HECTD3 (**Fig. 5b-f**). In all cases, we observed robust light-inducible knockdown, demonstrating the versatility and efficacy of OptoCas13d-on for controlling endogenous gene expression.

**Figure 5.**
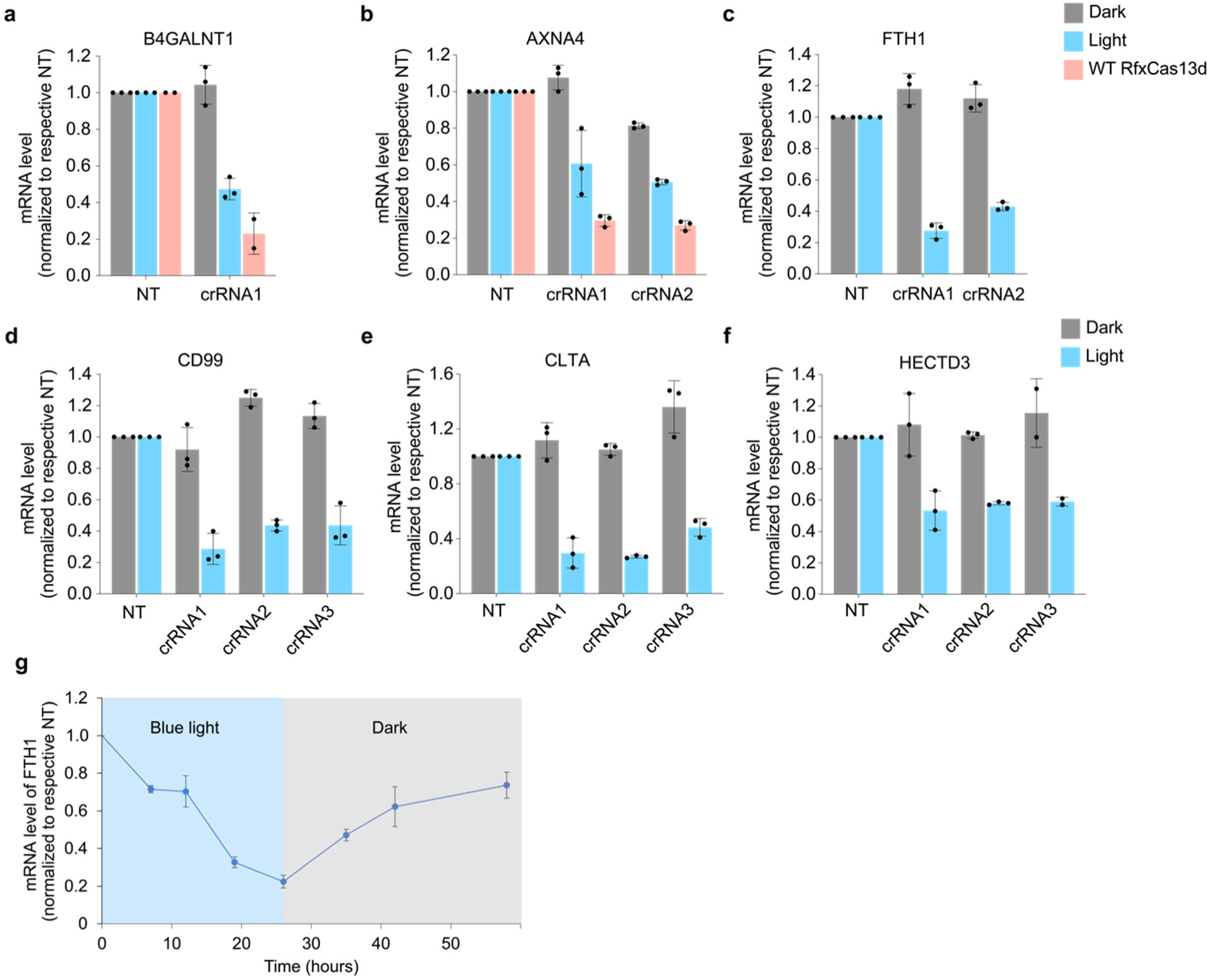
Endogenous transcript knockdown using OptoCas13d-on. (**a**) and (**b**) B4GALNT1 and AXNA4 mRNA level of RfxCas13d-LightR or wild type RfxCas13d clonal cell line transfected with B4GALNT1-targeting crRNA and AXNA4-targeting crRNA or non-targeting crRNA followed by incubation in dark or light condition. (**c**) - (**f**) Corresponding mRNA levels of RfxCas13d-LightR clonal cell line transfected with crRNA targeting different endogenous transcripts including AXNA4, FTH1, CD99, CLTA and HECTD3. All results are normalized to respective non-targeting crRNA groups. For B4GALNT1 crRNA, AXNA4 crRNA1, FTH1 crRNA1, CD99 crRNA1 and crRNA3, CLTA crRNA1, HECTD3 crRNA1 and crRNA3, biological replicates were performed; the rest of crRNA were carried out in technical replicates. (**g**) Time course experiment of RfxCas13d-LightR clonal cell line transfected with FTH1-targeting crRNA followed by 26 h light duration and then 30 h dark duration. mRNA levels were measured in some time points. Error bars indicate the standard deviation of n=3 technical replicates.

Finally, we determined the temporal dynamics and reversibility of light-induced knockdown after an acute change in illumination conditions. We conducted a time-course experiment using RfxCas13d-LightR transfected with an FTH1-targeting crRNA. Cells were illuminated for 26 hours, after which they were transferred to dark conditions for an additional 25 hours. mRNA levels were measured at multiple time points throughout the experiment (**Fig. 5g**). We observed a detectable reduction in FTH1 mRNA levels within 12 hours of light exposure, with a more pronounced knockdown as illumination duration increased, reaching a maximum effect between 18 and 26 hours. Upon transitioning cells to dark conditions at the 26-hour mark, FTH1 transcript levels gradually recovered over time, demonstrating the reversibility of the system. These results confirm that RfxCas13d-LightR provides a tunable and reversible platform for the controlled knockdown of endogenous genes with high temporal precision.

## Discussion

We have identified a single site in RfxCas13d (QK634) that allows for light and chemical induction *via* domain insertion. This modularity allowed us to develop OptoCas13d-off (active in dark and inactive in light), OptoCas13d-on (active in light and inactive in dark), and ChemoCas13d (active in presence of a chemical inducer). Recent studies have shown that toxicity associated with the promiscuous nuclease activity of Cas13 proteins can be mitigated by low effector and target RNA expression levels. Consistent with these findings, we observe selection against active Cas13 variants in our initial screen, and we find that light- and drug-switchable activity are optimal at specific RfxCas13d expression levels. Once appropriate expression levels are obtained, our OptoCas13d-on system exhibits high-quality stimulus-switchable control in stable cell line contexts.

We performed *in vitro* biochemical assays to gain insight into the switchable mechanism of OptoCas13d-off. We found that the insertion of AsLOV2 does not affect binding of OptoCas13d-off to its crRNA, and binding of the binary complex to its cognate target is only slightly impaired. In contrast, target RNA cleavage is substantially reduced in response to illumination. These data suggest that insertion of AsLOV2 at QK634 primarily affects RfxCas13d nuclease activity, not complex formation. Further validating these observations, we found that mammalian cells expressing a fusion of ADAR2 and a catalytically dead OptoCas13d-off exhibited similar levels of RNA editing in both light and dark, consistent with similar degrees of RNA complex formation in both conditions.

The idea of optogenetically and chemically inducible Cas13 proteins has been explored by others^13–15^. Notably, these systems were based on inducible dimerization to reconstitute split PspCas13b or RfxCas13d proteins to restore their activity, and none report light-controlled RNA degradation comparable to that achieved by full-length Cas13 enzymes. While powerful, split systems can result in leaky activity in the absence of induced dimerization as well as low enzyme activity after reconstitution. Our approach, identifying a domain insertion site for allosteric control, is quite effective in generating variants that exhibit either light or drug-induced enzymatic activity. This versatility provides end-users with the ability to knock down an RNA of interest either in the presence or absence of multiple stimulus modalities.

In the future, we envision that OptoCas13 could be used to control and perturb essential or tightly regulated cellular processes with high spatial and temporal precision, such as cell cycle progression, axonal RNA transport, and developmental gene expression. Such a tool could also be used to study biological processes such as investigating sub-populations of infected cells during viral-host interactions. The emergence of single-cell RNA sequencing has demonstrated that in many tissues and organisms, there are a diverse array of cell types, each with different gene regulatory networks. Our tool complements these sequencing efforts by enabling an end user to perturb RNA levels in a precisely defined subset of cells both in time and in space.

## Methods

### Plasmid construction

Plasmids associated with Mu transposition were originally obtained as a gift from David Savage which include pUCKanR-Mu-BsaI (Addgene #79769) and pATT-Dest (Addgene #79770)^23^. Coding sequence of RfxCas13d and expression vectors of RfxCas13d and crRNA were also obtained as a gift from Patrick Hsu which include pXR001: EF1a-CasRx-2A-EGFP (Addgene #109049) and pXR003: CasRx crRNA (Addgene #109053)^5^. pLZD047 was constructed by cloning RfxCas13d coding sequence from pXR001 into pATT-Dest with BsmBI restriction sites added on both side of RfxCas13d through in-fusion assembly (Takara #638911). Landing pad plasmids including recombination plasmids pJG_082 AttB_mCherry_Bgl, pJG_083 AttB_EGFP_Bgl2 and the pJG_075 Bxb1 which expresses Integrase essential for gene integration were obtained as gifts from Jacob Goell (Rice University, Hilton Lab). pLZD081 was constructed by cloning NLS-RfxCas13d-NLS into attB recombination plasmid through in-fusion assembly, and IRES-mCherry, the PCR product with pIRES2-mCherry-p53 deltaN (Addgene #49243) as template was tagged to NLS-RfxCas13d-NLS also through in-fusion assembly^29^. For transient transfection, Cas13d and its variants were cloned into PQ vector through in-fusion assembly, and crRNAs were cloned into pXR003. For stable expression, Cas13d and its variants tagged with IRES-mCherry were cloned into Piggybac vector downstream Tet-on promoter, and crRNAs were cloned into LRG2.1 vector which contains a constitutively expressed iRFP as the selection marker. LRG2.1 vector, which was originally LRG2.1-TagBFP2 (Addgene #124773) was obtained as a gift from Jason Sheltzer^30^. The AsLOV2 (408-543) sequence used in this study is the same as we previously used^22^. LightR sequence was cloned from LightR-bRaf-mVenus (Addgene #162154 ^16^). UniRapR sequence was cloned from pUSE-Src-YF-UniRapR-mCerulean-myc (Addgene #45381)^31^. See Supplementary Table 1 for a list of plasmids used in this study.

### Construction of AsLOV2 insertion library

To obtain a transposon that contains a chloramphenicol resistance gene, PCR was performed with pUCKanR-Mu-BsaI as template, 5′-taggcaccccaggctttacac-3′ as forward primer and 5′-tctgtaagcggatgccggga-3′ as reverse primer. The product was purified and digested with HindIII and BglII followed by purification with NucleoSpin Gel and PCR Clean-up Columns (Takara, #740609). The resulting DNA was directly used as transposon. Transposition reactions were conducted in a total volume of 20 μL with the following components: 0.14 pmol pLZD047, 0.55 pmol transposon DNA, 4 μL 5×MuA reaction buffer and 1 μL 0.22 μg/μL MuA transposase enzyme (Thermo Fisher Scientific, #F750). Reactions were incubated for 18 h at 30 °C followed by 10 min at 75 °C to heat inactivate MuA transposase. Completed reactions were cleaned up with DNA Clean & Concentrator-5 Kit (Zymo Research, #D4014) and eluted with 20 μL nuclease-free water (Thermo Fisher Scientific, #AM9932), which was then transformed into 200 μL TransforMax Electrocompetent E. coli (Lucigen, #EC10010). An aliquot of the recovery culture was spread on an LB agar plate with carbenicillin (Gold Biotechnology, #C-103-5) and chloramphenicol (Gold Biotechnology, #C-105-5) antibiotics to assess reaction efficiency. Remaining recovery culture was transferred to 50 ml LB with chloramphenicol and carbenicillin to select for plasmids with transposon insertion, and after overnight growth the library was collected with the Plasmid Plus Midi Kit (Qiagen, #12943), which was named pLZD047_CmR01. Qubit dsDNA Quantification Assay Kits (ThermoFisher Scientific, #Q32851) were used to quantify library DNA concentration in all the following steps.

To isolate library members containing RfxCas13d with CmR insertion, BsmBI digestion of pLZD047_CmR01 was performed. Total amount of 5 µg pLZD047_CmR01, 25 µL 10×NEBuffer 3.1, 50 units BsmBI_v2 (New England Biolabs, #R0739S) in a total volume of 250 µL was mixed and incubated 55°C for 1 h followed by 80°C for 20 min. Gel separation was performed afterwards and the band with the expected size of RfxCas13d with CmR insertion (4154 base pairs) was cut and purified with Zymoclean Gel DNA Recovery Kit (Zymo Research, #D4007), which was named pLZD047_CmR01_DNA. Next, to clone pLZD047_CmR01_DNA back to the original pLZD047 vector, backbone PCR was performed with pLZD047 as the template and primers as follows:

Forward primer: tttttcggtctccGGATCgagacggttaagatc
Reverse primer: tttttcggtctccGCTGGgagacgatggtatatctccttcttaaagttaaaca.

The PCR product was then digested with BsaI-HF_v2 (New England Biolabs, #R3733S) which created sticky ends on both sides that match up with the pLZD047_CmR01_DNA, and the digested product was termed pLZD047_DNA. T4 DNA ligase (New England Biolabs, #M0202S) was then used to ligate pLZD047_DNA with pLZD047_CmR01_DNA following the product protocol (https://www.neb.com/en-us/protocols/0001/01/01/dna-ligation-with-t4-dna-ligase-m0202), and the resulting library was termed pLZD047_CmR02.

Next, Golden Gate cloning was used to replace the chloramphenicol resistance gene with the AsLOV2 domain in pLZD047_CmR02. In detail, pLZ047_CmR02 was used as the backbone and insert was linear DNA with AsLOV2 (408–543) coding sequence and two BsaI digestion sites on both ends. Briefly, 75fmol of backbone and 210 fmol of AsLOV2 insert was mixed with 37.5 units BsaI_HFv2 (New England Biolabs, #R3733S), 2000 units T4 DNA Ligase (New England Biolabs, #M0202S) and 5 μL 10×T4 DNA Ligase Reaction Buffer in a total volume of 50 μL. The reaction was incubated 2 min at 37 °C, 5 min at 16 °C (first two steps cycled 50 times), 20 min at 60 °C and 20 min at 80 °C. Reactions were purified with DNA Clean & Concentrator-5 Kit and transformed into 100 μL TransforMax Electrocompetent *E. coli*. Cells were then transferred to 25 ml LB with carbenicillin, and the library was collected with the Plasmid Plus Midi Kit, which was named pLZD047_LOV. Another step of BsmBI digestion was performed for pLZD047_LOV and the band with size corresponding to RfxCas13d-LOV (3320 base pairs) was purified, with the name pLZD047_LOV_DNA

Lastly, Golden Gate cloning was used again to clone pLZD047_LOV_DNA into a landing pad recombination plasmid. PCR was performed with the pLZD81 as the template, forward primer taccatcgtcTCCGGATCCggacctaagaaaaagaggaaggtggcg and reverser primer cttaaccgtctcaGCTGGCCTCCACCTTtCTC to obtain the linearized landing pad recombination plasmid. For golden gate cloning, pLZD047_LOV_DNA was used as the insert and the linearized landing pad recombination plasmid was used as the backbone. Briefly, 58 fmol backbone and 78 fmol of insert were mixed with 15 units BsmBI_v2, 800 units T4 DNA Ligase and 2 μL 10×T4 DNA Ligase Reaction Buffer in a total volume of 20 μL. The reaction was incubated 2 min at 42 °C, 5 min at 16 °C (first two steps cycled 50 times), 20 min at 60 °C and 20 min at 80 °C. Reactions were purified using a DNA Clean & Concentrator-5 Kit and transformed into 100 μL TransforMax Electrocompetent *E. coli*. Cells were then transferred to 30 ml LB with carbenicillin, and the library was collected with the Plasmid Plus Midi Kit, which was named pLZD081_LOV.

### Construction of the reporter cell line for selections and transient transfection assay

HEK293T landing pad (293T-LP) cell line was a gift from Kenneth Matreyek (Case Western Reserve University), which was derived from HEK293T that had been bought from ThermoFisher. HEK293T landing pad cell line with constitutively-expressed eGFP (293T-LP-dGFP) was constructed by incorporating constitutively-expressed eGFP in pHR vector (pLZD075) into 293T-LP genome through lentivirus transduction. All the cells were kept in DMEM (ThermoFisher, #11995073) supplemented with 10% FBS (R&D Systems, #S11150), 1% Pen Strip (Gibco, #15140-122) and 2mM L-Glutamine (Gibco, #25030-081) throughout all experiments.

To produce lentiviral particles, HEK293T cells were plated on a 6-well plate grown up to 40% confluency. At that point they were co-transfected with pLZD075 and lentiviral packaging plasmids (pMD and CMV) with FuGENE HD (Promega, #E2311). Specifically, 1500 ng pLZD075, 1330 ng pCMVdR8.91 and 170 ng pMD2.G mixed with 9 μL Fugene HD transfection reagent were used for each well in a 6-well plate. Virus was collected after 53 h, filtered using a 0.45 mm filter. Polybrene (Sigma-Aldrich, #TR-1003-G) was added to the viral particles to the final concentration of 4 μg/mL. 293 T LP cell line was plated on a 6-well plate and infected with 350 μL of the virus at 40% confluency, and EGFP positive clonal cells were sorted 52 h post the infection time. Two weeks after clonal sorting, several clones were characterized through flow cytometry and one clone showing uniform EGFP expression was chosen as 293T-LP-dGFP for further incorporation of Cas13 variants or crRNA, or for transient transfection assays.

Next, EGFP-targeting crRNA (or non-targeting crRNA as a control) in LRG2.1 vector (pLZD078 for EGFP-targeting crRNA, and pLZD079 for non-targeting control crRNA) was further incorporated into 293T-LP-dGFP through lentivirus transduction, with a constitutively-expressed iRFP as the selection marker. To produce lentiviral particles, HEK 293 T cells were plated on a 6-well plate grown up to 40% confluency. At that point they were co-transfected with pLZD078 or pLZD079 and lentiviral packaging plasmids (pMD and psPAX2.0) with FuGENE HD (Promega, #E2311). Specifically, 1500 ng pLZD078, 1850 ng pCMVdR8.91 and 300 ng pMD2.G mixed with 10.95 μL Fugene HD transfection reagent were used for each well in a 6-well plate. Virus was collected after 50 h, filtered using a 0.45 mm filter. Polybrene (Sigma-Aldrich, #TR-1003-G) was added to the viral particles to the final concentration of 4 μg/mL. 293 T LP cell line was plated on a 6-well plate and infected with 300 μL of the virus at 40% confluency, and iRFP clonal cell was sorted 48 h post the infection time. Two weeks after clonal sorting, several clones were characterized through flow cytometry and one clone showing uniform iRFP expression was chosen as the reporter cell line for following negative selections experiments, termed 293T-LP-dGFP-GFP_crRNA or 293T-LP-dGFP-NT_crRNA.

### Integration of Cas13-LOV insertion library into 293T landing pad reporter cell line and follow-up negative selections

HEK293T-LP-dGFP_crRNA cell lines were plated on 6-well plates one day before transfection. Recombination was performed by transfecting cells with 1500 ng of pJG75 Bxb1 and 2500 ng of AsLOV2 insertion library pLZD081_LOV in doxycycline-free media with 9 μL FuGENE HD transfection reagent. 32 h following transfection, the media was changed to media supplemented with 2 μg/mL doxycycline. 62h after media replacement cells that are mCherry positive and BFP negative were sorted to a new 6-well plate with flow cytometry. Before flow cytometry, cells were detached with trypsin and resuspended in DMEM containing 10% serum. Flow cytometry was performed with SH800S Cell Sorter equipped with Sony 100 μm Sorting Chip (Sony Biotechnology, LE-C3210). BFP was excited with a 405 nm laser, and emitted light was collected after passing through 450/50 nm band pass filters. EGFP was excited with a 488 nm laser, and emitted light was collected after passing through 525/50 nm bandpass filters. mCherry was excited with a 561 nm laser, and emitted light was collected after passing through 665/30 nm band pass filters. Manual compensation was performed to decouple crossovers among BFP, EGFP and mCherry signals Before sorting, live, single cells were gated using FSC-A and SSC-A (for live cells) and FSC-A and FSC-H (for single cells) and at least 100,000 cells were sorted and plated on a new 6-well plate in doxycycline-free media. Another round of doxycycline induction and sorting was performed two weeks after the first sorting to further enrich mCherry-positive and BFP-negative cells.

### Extraction of genomic DNA and next-generation sequencing

PureLink Genomic DNA Mini kit (Invitrogen, #K182001) was used to extract genomes from library cells following the protocol in manual, and PCR was performed with forward primer 5′-CCAGGGCTCGAGACCGCAACTACACGCCACC-3′ and reverse primer 5′-AGCTTCGAATTCGGGGCGGATCAGCTTGGTAC-3′ to amplify the library fragments from the genome. Library fragment DNA was then sheared with a Covaris S220 focused ultrasonicator using AFA microTUBEs (Covaris, #PN 520052) to an approximate size of 300–400 base pairs. NEBNext® Ultra™ II DNA Library Prep with Sample Purification Beads (New England Biolabs, #E7103S) was used to prepare DNA library for sequencing from sheared DNA. Sheared DNA and prepared samples were analyzed for size distribution on an Agilent 2100 Bioanalyzer using DNA 1000 chips (Agilent Technologies). Double-stranded DNA concentrations of the adaptor-prepared samples were measured with a dsDNA HS Assay Kit (Invitrogen, #Q32851) on a Qubit Fluorometer. A normalized pool of samples was run on a MiSeq Nano 300 nt or MiSeq Micro 300 nt for over 300 cycles. Analysis of FASTQ files of sequencing results was performed through MATLAB R2021b, (with the script provided in Supplementary Note 4). For deeper sequencing to map all of the initial integration sites in Figure 1D, Nanopore sequencing of the same initial libraries was carried out by Plasmidsaurus and the AsLOV2 insertion sites into Cas13 were mapped using custom Rust code. All code used to map reads is available on the Toettcher laboratory’s Github page: https://github.com/toettchlab/Zhu-Nguyen2025.

### Cell transient transfection for EGFP knockdown assay

In all Cas13 variants transient transfection for EGFP-knock down assays, plasmid encoding Cas13 variant and plasmid encoding crRNA were double transfected to 293T-LP-dGFP. For transfection, Cas13d variant coding sequence is cloned to the PQ vector, and EGFP targeting crRNA (GFP_crRNA) or non-targeting control crRNA (NT_crRNA) was cloned into the pXR003 vector. 293T-LP-dGFP cells were plated on 12-well plates 1 day prior to transfection. 400 ng of Cas13 plasmid and 400 ng of crRNA plasmid were mixed with 2.4 μL FuGENE HD Transfection Reagent and transfected to 293T-LP-dGFP. For light-based assay, All the cells were kept in the dark for 5 h post transfection and then irradiated with 450 nm blue LEDs at an intensity of 0.80 mW/cm^2^ or remained in the dark for additional 26 h before characterization with flow cytometry. For rapamycin-based assay, 1 h prior to transfection cell media was replaced with fresh media with 0.01 μM rapamycin (Sigma-Aldrich, #553211), and flow cytometry was performed 28 h after transfection.

### Protein expression and purification

Protein expression plasmid was constructed by cloning a mammalian codon-optimized CasRx gene (gift from Patrick Hsu, Addgene #109049 and #109050 for catalytically active and inactivate variants) into a bacterial expression vector bearing 6xHis and TEV cleavage sites (gift from Scott Gradia, Addgene #29653). AsLOV2 domain was then inserted into this expression backbone using Infusion cloning.

For protein expression, plasmids were transformed into Rosetta™ 2(DE3) Singles™ competent cells following the manufacturer’s instructions. Individual colonies were picked the next day and inoculated in 25-30 mL LB broth for 14-18 h. The culture was then scaled up to 2 L with Terrific Broth and continued to be shaked at 37C until OD600 = 0.8-1.0. The culture was next placed on ice for 30-45 minutes followed by the addition of 1 mM IPTG to induce CasRx expression. The culture was transferred to a shaker and incubated at 18C for 14-18 h.

For protein purification, cell pellets were harvested via centrifugation. The pellets were then resuspended in lysis buffer (500 mM NaCl, 50 mM Tris-HCl pH = 7.5, 20 mM imidazole, 1 mM TCEP-HCl, 0.5 mg/mL lysozyme, 0.5 mM PMSF, and 0.1 mg/mL DNase I) and subjected to sonication. Cell lysate was then centrifuged at 40,000 ×g for 30 min followed by filtering through 0.22 um syringe filter. The clarified lysate was injected into Nuvia IMAC, Ni-charged (Biorad) pre-equilibrated with buffer A (500 mM NaCl, 50 mM Tris-HCl pH = 7.5, 20 mM imidazole, 1 mM TCEP-HCl) via the NGC Quest 10 Plus FPLC system (Biorad, #12009287). In the case of OptoCas and dOptoCas, 5mM Flavin Mononucleotide was added to buffer A prior to purification The column was eluted using buffer B (500 mM NaCl, 50 mM Tris-HCl pH = 7.5, 300 mM imidazole, 1 mM TCEP-HCl), pooled together and dialyzed against buffer C (250 mM NaCl, 50 mM Tris-HCl pH = 7.5, 300 mM imidazole, 1 mM TCEP-HCl, 10% glycerol) overnight. The next day, the protein mixture was concentrated and injected into the cation exchange column Macro-Prep High S (Biorad, #12009272) using the same FPLC system. The eluted fractions were collected by gradually exchanging buffer C with buffer D (2 M NaCl, 50 mM Tris-HCl pH = 7.5, 300 mM imidazole, 1 mM TCEP-HCl, 10% glycerol) in a gradient fashion. Protein purity was analyzed using gel electrophoresis, pooled together, dialyzed against final buffer (600 mM NaCl, 50 mM Tris-HCl pH = 7.5, 300 mM imidazole, 2 mM DTT, 10% glycerol), flash frozen and stored at −80C until use.

### In vitro trans-cleavage assay

Cleavage assays were adapted from previously described protocols^27^. A master mix was created with 1× reaction buffer (20 mM HEPES pH 8.0 with 60 mM KCl and 3.5% PEG-8000), 10 mM MgOAc, 1.33 U µl−1 murine RNase inhibitor (New England Biolabs), 6.25 μM of crRNA (IDT), and 5 μM of either WT Cas13d or Cas13d containing AsLOV2. The crRNA and Cas13d incubated at 37°C for 15 minutes to complex under blue light or in the dark. Subsequently, 50 ng of target RNA was added to the reaction and incubated at 37°C for another 15 minutes. Water was used in replacement of target for no target controls (NTCs). Subsequently, 133.33 nM polyU (6 uracils; IDT)) quenched HEX reporter was added to each reaction. 15 μL reactions were loaded in technical triplicate onto a Greiner 384 well clear-bottom microplate (item no. 788096). The reactions were incubated at 37 °C for up to 3 hours with fluorescent readings taken every 5 minutes using an Agilent BioTek Cytation 5 microplate reader (excitation: 485 nm, emission: 525 nm).

### Electrophoretic Mobility Shift Assay (EMSA)

For in vitro gel shift assays to visualize binary complex formation, 20 μL samples were prepared to reach final concentrations of 500 nM 3’FAM-GFP targeting-crRNA and either 400 nM wild-type CasRx or 400 nM CasRx-AsLOV2 in 1× binding buffer (24 mM KCl, 4mM Tris-HCl, pH8.0, 0.4 mM DTT, 10% glycerol, 0.1 mg/mL BSA, 5mM MgCl_2_). For samples meant to visualize ternary complex formation, 80 nM Cy5-labeled EGFP target RNA was also included. After preparation, samples were allowed to complex either in the dark or under blue light for 15 minutes at 37C before being mixed with gel loading dye and run on 5% polyacrylamide gels (Biorad, #4565013) in 0.5× TBE buffer. Gel boxes were placed on ice and kept under aluminum foil to prevent light interference following complexing. crRNA signal was visualized by imaging the FAM channel on an Azure Biosystems 600 imager. Gels were then stained with SYBR gold (Invitrogen, #S11494) and re-imaged using the SYBR gold channel in order to visualize both crRNA and target RNA.

### Integration of Cas13d variants into 293T-LP-dGFP with Piggybac transposition

RfxCas13d variants coding sequences tagged with IRES-mCherry were first cloned to Piggybac vector downstream the tetracycline (Tet) inducible promoter. One example is pLZD239 which contains Cas13LightRQK634-IRES-mCherry. rtTA (reverse tetracycline-controlled transactivator) was already stably expressed in 293T-LP-dGFP cell lines. To incorporate pLZD239 into 293T-LP-dGFP (or 293T-LP-dGFP-GFP_crRNA with the same protocol), 2000ng pLZD239, 500ng Piggybac helper plasmid was mixed with 18 μL FuGENE HD Transfection Reagent and transfected to 293T-LP-dGFP. The cells were maintained in doxycycline-free media unless otherwise mentioned. Six days after transfection, the media was changed to media supplemented with 2 μg/mL doxycycline. After another 56h, flow cytometry was performed and mCherry positive cells were bulk sorted. One week afterwards another round of flow cytometry was performed, and individual mCherry-positive cells were sorted into each well of a 96-well plate. The resulting clonal cell line was termed 293T-LP-dGFP-Cas13LightR or 293T-LP-dGFP-GFP_crRNA-Cas13LightR.

### Base editing of Luciferase Reporter

Dual secreted luciferase reporter (EF1alpha-Gaussia luciferase, CMV-Cypridina luciferase) was obtained as a gift from Feng Zhang (Addgene plasmid # 181934) and modified to install a premature stop codon at amino acid 85 of CLuc gene. 18-24 hours prior to transfection, 10,000 HEK293T cells were seeded in two 96-well plates. Next, 40 ng of dual luciferase reporter plasmid, 150 ng of wild-type dRfxCas13d-ADAR or dOptoCas13d-QK634-AsLOV2-ADAR protein, and 300 ng of crRNA targeting cLuc were mixed with Lipofectamine 3000 and P3000 reagent (Invitrogen, #L3000015) and co-transfected into each well. Four hours later, one of the 96-well plates was then treated under blue light for 48 hours. Medium containing luciferase in both light and dark treated plates were used for luciferase analysis using UltraBriteTM Cypridina-Gaussia dual luciferase assay reagent (Targeting systems, #DLAR-4 SG-1000) and Biotek Cytation 5 plate reader with automatic gain and 1 sec exposure time (Agilent). Due to constitutive expression of GLuc, its signal was adjusted 100-fold lower than the actual measurements prior to background normalization with CLuc signal.

### Cell transient transfection for knocking down endogenous transcripts

For transfection, crRNA targeting endogenous transcripts or non-targeting control crRNA (NT_crRNA) was cloned into pXR003 vector. 293T-LP-dGFP-Cas13LightR was used which contains doxycycline-inducible Cas13LightRQK634. Cells were plated on 12-well plates in 2 μg/mL doxycycline media 1 day prior to transfection. During transfection, 800 ng of crRNA plasmid were mixed with 2.4 μL FuGENE HD Transfection Reagent and transfected to 293T-LP-293T-LP-dGFP-Cas13LightR. All the cells were kept in the dark for 5 h post transfection and then irradiated with 450 nm 5 mm blue LEDs (Digikey, Inc) at an intensity of 0.80 mW/cm^2^ or remained in the dark for additional 32 h before follow-up RT-qPCR experiments.

### RT-qPCR

To analyze relative expression of endogenous transcripts, total RNA was extracted from cells with RNease Mini Kit (Qiagen #74106) following protocol in the manual. Reverse transcription was then performed with LunaScript RT SuperMix Kit (NEB #E3010), and the products were directly mixed with qPCR primers and Luna Universal qPCR Master Mix (NEB #M3003L) for qPCR assay. The experiment follows the protocol for Two-step RT-qPCR using the LunaScript RT SuperMix Kit (NEB #E3010) and the Luna Universal qPCR Master Mix (NEB #M3003) available in the NEB website. Sequences of crRNA and qPCR primers are provided in Supplementary Table 2 and Supplementary Table 3.

## Author Contributions

Conceptualization, L.Z., L.T.N., C.M., J.E.T.; Methodology, L.Z., L.T.N., H.O., J.E.T.; Investigation, L.Z., L.T.N., A.G.B., K.M.G., J.H..; Funding, L.T.N., C.M., J.E.T.; Writing and Editing, L.Z., L.T.N., C.M., J.E.T.; Supervision, C.M., J.E.T.

## Acknowledgements

We thank all members of the Toettcher and Myhrvold labs for helpful discussions. This work was supported by the Omenn-Darling Bioengineering Institute (to L.T.N.), the National Institutes of Health grants R01GM144362 and U01DK127429 (to J.E.T.), R01AI182281 (to C.M.), and T32GM007388 and T32GM148739 (to A.G.B. and K.M.G.), as well as a Princeton AI Lab Seed Grant 2025-74 (to C.M. and J.E.T.).

