## Supplementary Information for "Multimodal control of Cas13 activity through domain insertion at an allosteric hotspot"

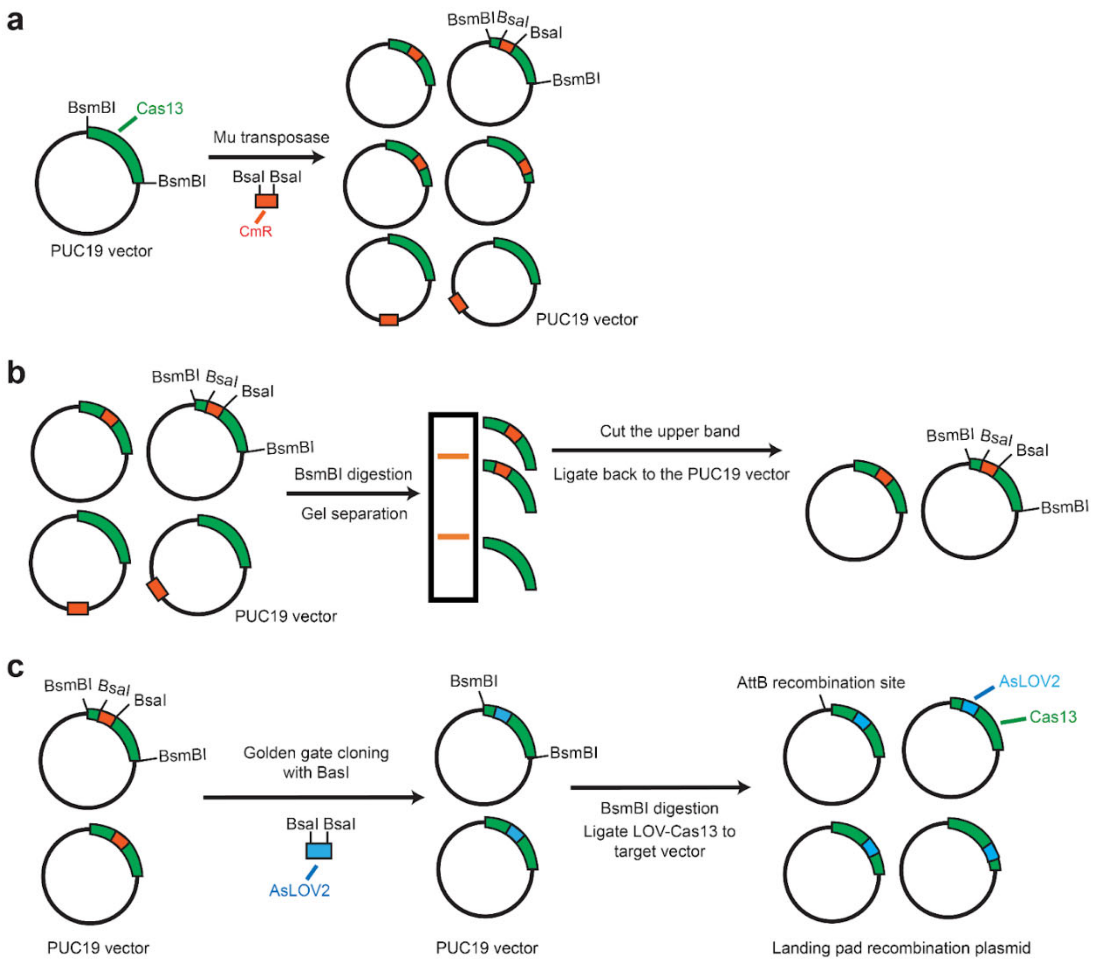


**Figure S1. Schematic of generation of OptoCas13d variant library. (a)** Transposition reaction to randomly insert chloramphenicol-resistant genes (CmR) into a pUC19 vector containing RfxCas13d coding sequences. **(b)** BsmBI digestion and gel separation to purify fragments of RfxCas13d with CmR insertion and ligate them back to the original PUC19 vector with two BsmBI restriction sites on both sides of RfxCas13d sequence present. **(c)** BsaI-based golden gate cloning to replace the CmR cassette with AsLOV2 domain, followed by BsmBI digestion and ligation of RfxCas13d-LOV fragments to the landing pad recombination plasmid for follow-up selections.

a

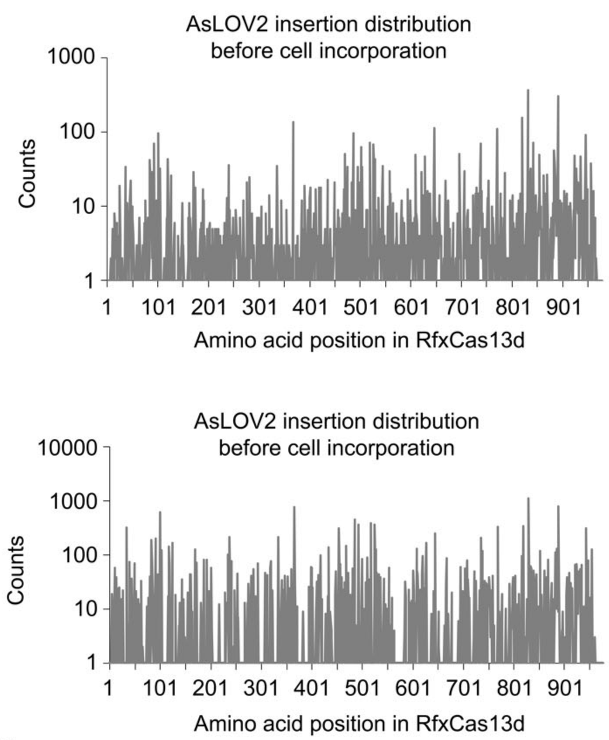


b

| Insertion site | Percentages before integration | Percentages after integration |
|---|---|---|
| 47 | 0.0038 | 0.0001 |
| 88 | 0.0025 | 0.0001 |
| 90 | 0.0050 | 0 |
| 191 | 0.0020 | 0 |
| 269 | 0.0020 | 0 |
| 391 | 0.0032 | 0 |
| 449 | 0.0036 | 0.0010 |
| 469 | 0.0021 | 0 |
| 498 | 0.0036 | 0 |
| 505 | 0.0059 | 0.0005 |
| 559 | 0.0052 | 0.0002 |
| 573 | 0.0013 | 0.0000 |
| 581 | 0.0007 | 0 |
| 582 | 0.0020 | 0 |
| 634 | 0.0027 | 0 |
| 637 | 0.0025 | 0 |
| 639 | 0.0025 | 0.0004 |
| 657 | 0.0023 | 0 |
| 674 | 0.0020 | 0 |
| 731 | 0.0030 | 0.0004 |
| 777 | 0.0025 | 0 |
| 786 | 0.0048 | 0.0005 |
| 797 | 0.0020 | 0.0002 |
| 817 | 0.0025 | 0 |
| 898 | 0.0018 | 0.0001 |
| 903 | 0.0027 | 0.0001 |
| 961 | 0.0027 | 0 |

c

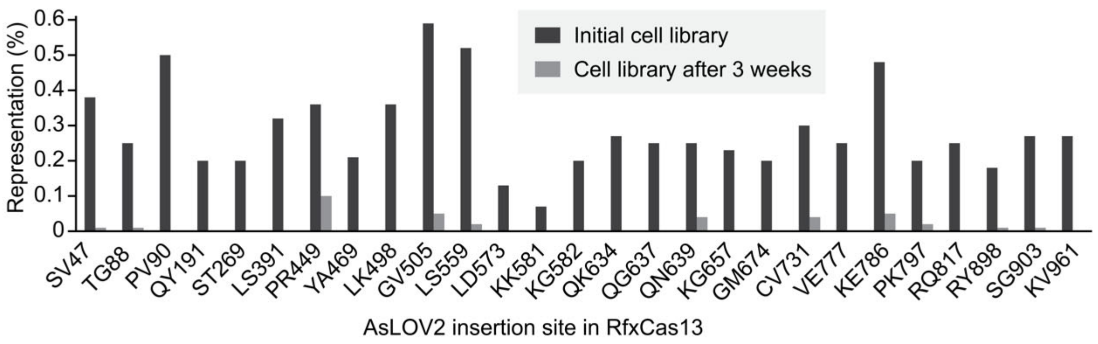


**Figure S2. Loss of some OptoCas13d library before and after cell incorporation reveals potential switchable variants.** (**a**) AsLOV2 insertion distribution of RfxCas13-LOV2 library before and after cell integration as determined by next-generation sequencing. (**b**) Insertion sites disappeared after integrating the RfxCas13d-LOV2 library into the landing pad reporter cell line. (**c**) Negative selection screening identifies 27 insertion sites that might be active variants.

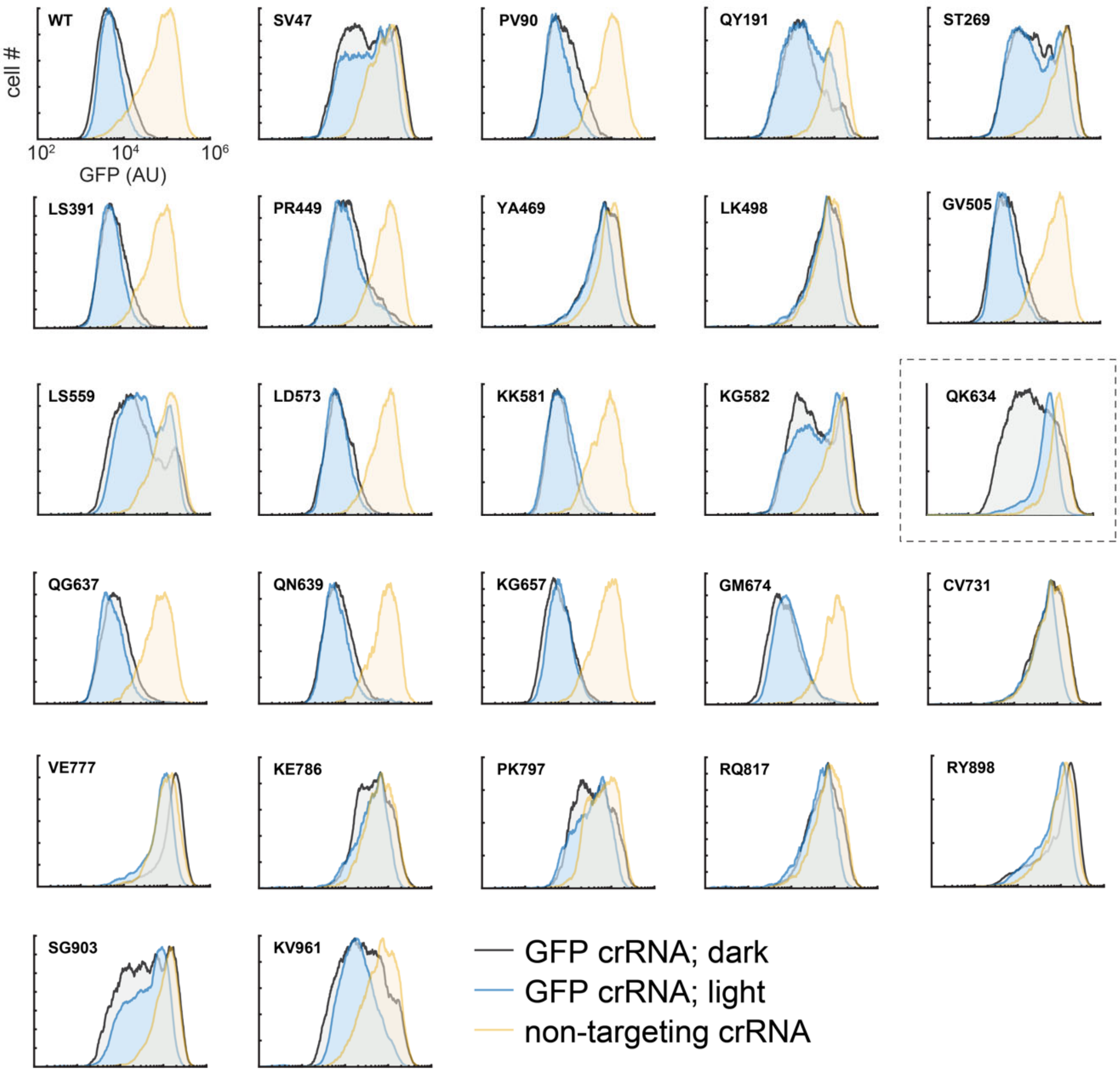


**Figure S3. Results of 26 candidates being tested.** EGFP stably expressing cells were transfected with RfxCas13d with AsLOV2 insertion at different sites followed by incubation in light or dark conditions and flow cytometry analysis.

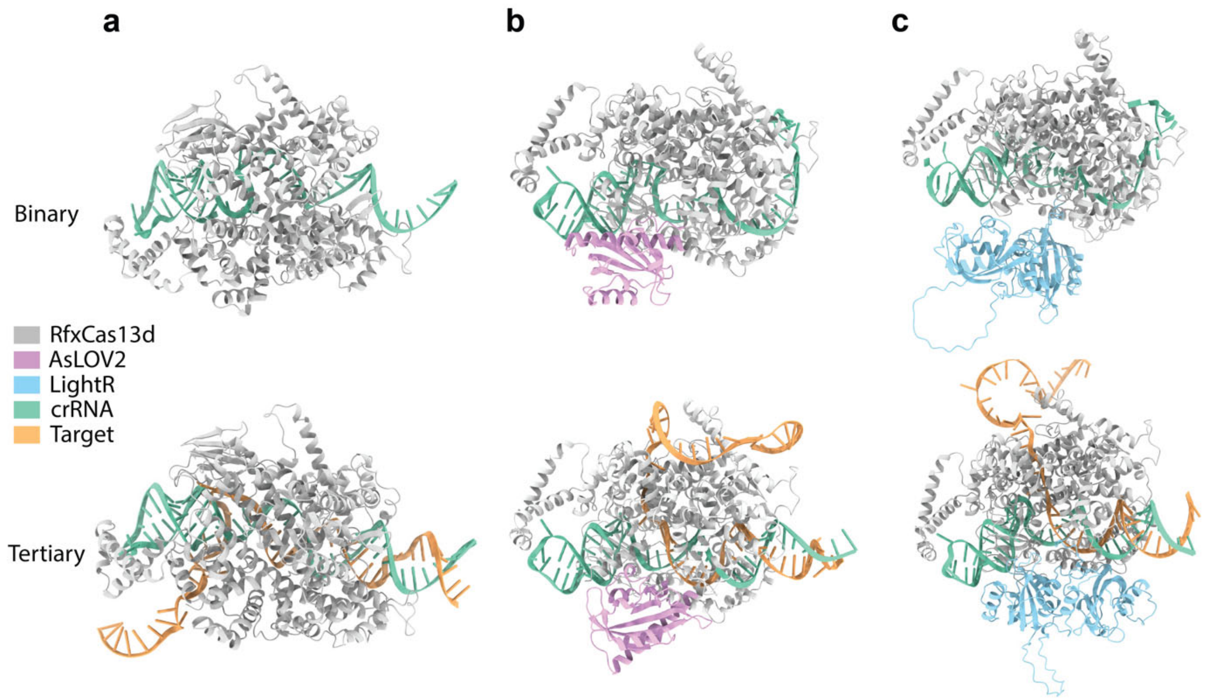


**Figure S4**. **Predicted structure of binary and tertiary complexes of RfxCas13d with various inserts**. (**a**) wild-type RfxCas13d. (**b**) RfxCas13d-AsLOV2. (**c)** RfxCas13d-LightR. crRNA contains a spacer targeting EGFP mRNA. Prediction was performed using AlphaFold 3.

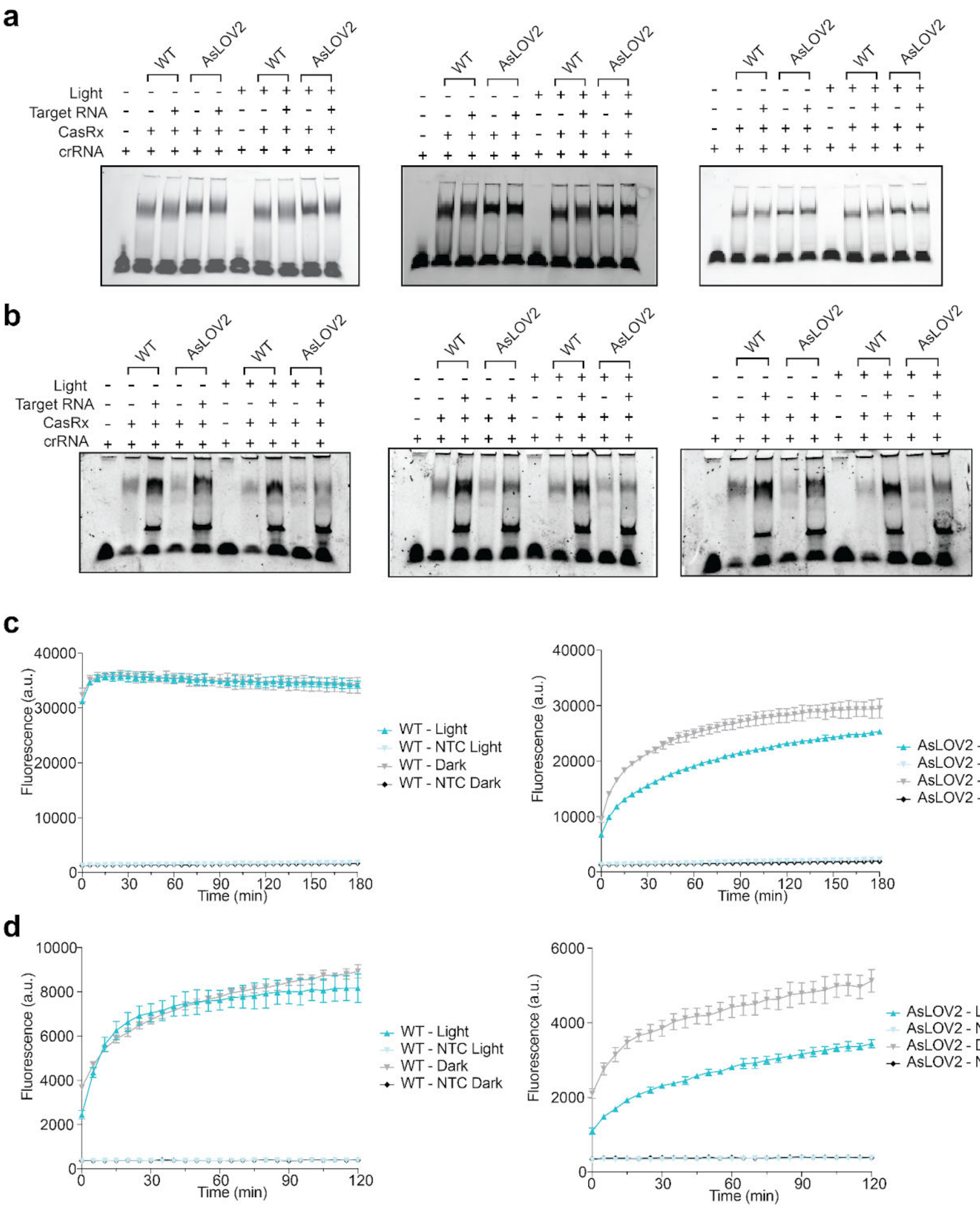


**Figure S5. *In Vitro* Characterization of Optocas13d-off and wild-type RfxCas13d**. (**a**) All 3 replicates for data shown in Figure 3d. (**b**) All 3 replicates for data shown in Figure 3f. (**c**) Cleavage kinetics for WT (left) and OptoCas13d-off (right) at 126.67 nM of crRNA and 100 nM of effector. (**d**) Cleavage kinetics for WT (left) and OptoCas13d-off (right) at 6.33 nM of crRNA and 5 nM of effector. Error bars in **c-d** show mean + SD for 3 independent replicates.

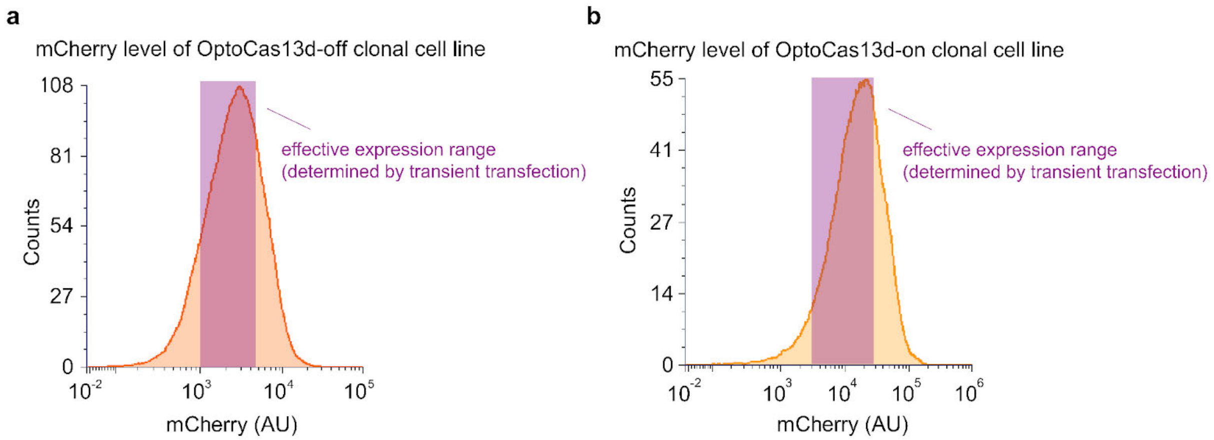


**Figure S6. Effective range of OptoCas13d-off and OptoCas13d-on expression levels that allow for robust light switchable activity.** (**a**) mCherry histogram of OptoCas13d-off clonal cell line. (**b**) mCherry histogram of OptoCas13d-on clonal cell line. Both OptoCas13d-off and OptoCas13d-on were tagged with IRES-mCherry.

**Supplementary Tables**

**Supplementary Table 1: Plasmids used for this study**

| Plasmids | Insert1 | Insert2 | Vector | Sources |
|---|---|---|---|---|
| pUCKanR-Mu-BsaI | Transposon-propagation vector used for Mu-BsaI modified transposon | | | Addgene #79769 |
| pATT-Dest | Vector used for transposition | | | Addgene #79770 |
| pXR001: EF1a-CasRx-2A-EGFP | NLS-RfxCas13d-NLS-P2A-EGFP | | pXR001 | Addgene #109049 |
| pIRES2-mCherry-p53 deltaN | p53 deltaN | mCherry | | Addgene #49243 |
| LRG2.1-TagBFP2 | guide RNA | TagBFP2 | | Addgene #124773 |
| LightR-bRaf-mVenus | LightR-bRaf-mVenus | | | Addgene #162154 |
| pUSE-Src-YF-UniRapR-mCerulean--myc | UniRapR-mCerulean | | | Addgene #45381 |
| pLZD3 | Non-targeting gRNA | | pxr003 | Addgene #109053 |
| pLZD4 | GFP-targeting gRNA | | pxr003 | This study |
| pLZD47 | CasRx | | PUC19 | This study |
| pLZD78 | U6_GFP-targeting gRNA | EF1a-iRFP | LRG2.1 | This study |
| pLZD79 | U6_none-targeting gRNA | EF1a-iRFP | LRG2.1 | This study |
| pLZD81 | CasRx-IRES-mCherry | | AttB recombination | This study |
| pLZD151 | CasRx-AsLOV2(QK634)-IRES-mCherry | | AttB recombination | This study |
| pLZD182 | EF1a_CasRx-IRES-mCherry | | PQ | This study |
| pLZD183 | EF1a_CasRx-AsLOV2(QK634)-IRES-mCherry | | PQ | This study |
| pLZD185 | CasRx_LightR(QK634)-IRES-mCherry | | PQ | This study |
| pLZD189 | B4GALNT1-targeting gRNA | | pxr003 | This study |
| pLZD222 | AXNA4-targeting gRNA | | pxr003 | This study |
| pLZD224 | AXNA4-targeting gRNA (II) | | pxr003 | This study |

| | | | | |
|---|---|---|---|---|
| pLZD239 | TREp_CasRx-LightR(QK634)-IRES-mCherry | | Piggybac | This study |
| pLZD240 | TREp_CasRx-IRES-mCherry | | Piggybac | This study |
| pLZD241 | TREp_CasRx-AsLOV2(QK634)-IRES-mCherry | | Piggybac | This study |
| pLZD247 | FTH1-targeting gRNA (II) | | pxr003 | This study |
| pLZD248 | FTH1-targeting gRNA | | pxr003 | This study |
| pLZD249 | CD99-targeting gRNA (II) | | pxr003 | This study |
| pLZD250 | CD99-targeting gRNA | | pxr003 | This study |
| pLZD251 | CD99-targeting gRNA (III) | | pxr003 | This study |
| pLZD252 | CLTA-targeting gRNA (II) | | pxr003 | This study |
| pLZD253 | CLTA-targeting gRNA | | pxr003 | This study |
| pLZD254 | CLTA-targeting gRNA (III) | | pxr003 | This study |
| pLZD255 | HECTD3-targeting gRNA | | pxr003 | This study |
| pLZD256 | HECTD3-targeting gRNA (II) | | pxr003 | This study |
| pLZD257 | HECTD3-targeting gRNA (III) | | pxr003 | This study |

#### Supplementary Table 2: gRNA sequences

| Plasmid | Description | gRNA spacer (5' to 3') |
|---|---|---|
| pLZD3 | Non-targeting gRNA | GCTACGTGGCTTCGCCTGTCGC |
| pLZD4 | GFP-targeting gRNA | CTTCACCTCGGCGCGGGTCTTG |
| pLZD189 | B4GALNT1-targeting gRNA | CCTCCTGACCAGAAGCTGCCTG |
| pLZD222 | AXNA4-targeting gRNA | CTTGTAGGCTGTCCTGATCTCC |
| pLZD224 | AXNA4-targeting gRNA (II) | GAATGTCATCTTCAAGGCTCCG |
| pLZD247 | FTH1-targeting gRNA (II) | GCTCTCCCAGTCATCACAGTCTGGTTTCTT |
| pLZD248 | FTH1-targeting gRNA | CTGAAGGAAGATTCGGCCACCTCGTTGGTT |
| pLZD249 | CD99-targeting gRNA (II) | AACGCCATCCGCAAGGTCAGCATCTGAAAA |
| pLZD250 | CD99-targeting gRNA | GTCAGCATCTGAAAAGCTACCGGAGGAACT |
| pLZD251 | CD99-targeting gRNA (III) | ATGGCTCCAGCCACGGCGACCACGACAGCC |
| pLZD252 | CLTA-targeting gRNA (II) | GGAGATGAGGACTGAGCGCATGCGGGAGAC |
| pLZD253 | CLTA-targeting gRNA | TCCACTCTGCTTCTTGCTTCCGAGAATTGG |
| pLZD254 | CLTA-targeting gRNA (III) | GTGGCTCTTCAGTGCACCAGCGGGGCCTGC |
| pLZD255 | HECTD3-targeting gRNA | TGGTATCCACTGTGAGTAGCAGCTTCTTGA |
| pLZD256 | HECTD3-targeting gRNA (II) | GCTCTCCACATACTGCTTCACGCTACCCAG |
| pLZD257 | HECTD3-targeting gRNA (III) | GCGTCTGTGGTCTCGTAGCCCAGCTTGTCT |

#### Supplementary Table 3: qPCR primers

| Transcripts | Forward primer (5'-3') | Reverse primer (5'-3') |
|---|---|---|
| GAPDH | ACAACTTTGGTATCGTGGAAGG | GCCATCACGCCACAGTTTC |
| GFP | AAGCAGAAGAACGGCATCAA | GGGGGTGTTCTGCTGGTAGT |
| B4GALNT1 | TGAGGCTGCTTTCACTATCCGC | GAGGAAGGTCTTGGTGGCAATC |
| AXNA4 | GATGCTCTCGTGAGACAGGATG | GCAACAGGTGATTTCGGTTCCG |
| FTH1 | TGAAGCTGCAGAACCAACGAGG | GCACACTCCATTGCATTCAGCC |
| CD99 | GAAAAGGAGGCAGTGATGGTGG | TGAAGCTAGAGATGGCTCCAGC |
| CLTA | CGATTGCAGTCAGAGCCTGAAAG | TAGCTGCTCGTCCTGTCTTGCA |
| HECTD3 | ATCGAGATCCGCATCGTGGAGT | TAGTTGGCTGGAACAGGTCTGC |

### Supplementary Notes

#### Supplementary Note 1: sequence of RfxCas13d-AsLOV2(QK634)

In black: RfxCas13d In green: Linker In blue: AsLOV2 (408-543)

ATCGAAAAAAAAAAGTCCTTCGCCAAGGGCATGGGCGTGAAGTCCACACTCGTGTCCGGCTCCAAA
GTGTACATGACAACCTTCGCCGAAGGCAGCGACGCCAGGCTGGAAAAGATCGTGGAGGGCGACAG
CATCAGGAGCGTGAATGAGGGCGAGGCCTTCAGCGCTGAAATGGCCGATAAAAACGCCGGCTATAA
GATCGGCAACGCCAAATTCAGCCATCCTAAGGGCTACGCCGTGGTGGCTAACAACCCTCTGTATACA
GGACCCGTCCAGCAGGATATGCTCGGCCTGAAGGAAACTCTGGAAAAGAGGTACTTCGGCGAGAGC
GCTGATGGCAATGACAATATTTGTATCCAGGTGATCCATAACATCCTGGACATTGAAAAAATCCTCGC
CGAATACATTACCAACGCCGCCTACGCCGTCAACAATATCTCCGGCCTGGATAAGGACATTATTGGA
TTCGGCAAGTTCTCCACAGTGTATACCTACGACGAATTCAAAGACCCCGAGCACCATAGGGCCGCTT
TCAACAATAACGATAAGCTCATCAACGCCATCAAGGCCCAGTATGACGAGTTCGACAACTTCCTCGA
TAACCCCAGACTCGGCTATTTCGGCCAGGCCTTTTTCAGCAAGGAGGGCAGAAATTACATCATCAAT
TACGGCAACGAATGCTATGACATTCTGGCCCTCCTGAGCGGACTGAGGCACTGGGTGGTCCATAAC
AACGAAGAAGAGTCCAGGATCTCCAGGACCTGGCTCTACAACCTCGATAAGAACCTCGACAACGAAT
ACATCTCCACCCTCAACTACCTCTACGACAGGATCACCAATGAGCTGACCAACTCCTTCTCCAAGAA
CTCCGCCGCCAACGTGAACTATATTGCCGAAACTCTGGGAATCAACCCTGCCGAATTCGCCGAACAA
TATTTCAGATTCAGCATTATGAAAGAGCAGAAAAACCTCGGATTCAATATCACCAAGCTCAGGGAAGT
GATGCTGGACAGGAAGGATATGTCCGAGATCAGGAAAAATCATAAGGTGTTCGACTCCATCAGGACC
AAGGTCTACACCATGATGGACTTTGTGATTTATAGGTATTACATCGAAGAGGATGCCAAGGTGGCTG
CCGCCAATAAGTCCCTCCCCGATAATGAGAAGTCCCTGAGCGAGAAGGATATCTTTGTGATTAACCT
GAGGGGCTCCTTCAACGACGACCAGAAGGATGCCCTCTACTACGATGAAGCTAATAGAATTTGGAGA
AAGCTCGAAAATATCATGCACAACATCAAGGAATTTAGGGGAAACAAGACAAGAGAGTATAAGAAGA
AGGACGCCCCTAGACTGCCCAGAATCCTGCCCGCTGGCCGTGATGTTTCCGCCTTCAGCAAACTCA
TGTATGCCCTGACCATGTTCCTGGATGGCAAGGAGATCAACGACCTCCTGACCACCCTGATTAATAA
ATTCGATAACATCCAGAGCTTCCTGAAGGTGATGCCTCTCATCGGAGTCAACGCTAAGTTCGTGGAG
GAATACGCCTTTTTCAAAGACTCCGCCAAGATCGCCGATGAGCTGAGGCTGATCAAGTCCTTCGCTA
GAATGGGAGAACCTATTGCCGATGCCAGGAGGGCCATGTATATCGACGCCATCCGTATTTTAGGAAC
CAACCTGTCCTATGATGAGCTCAAGGCCCTCGCCGACACCTTTTCCCTGGACGAGAACGGAAACAA
GCTCAAGAAAGGCAAGCACGGCATGAGAAATTTCATTATTAATAACGTGATCAGCAATAAAAGGTTCC
ACTACCTGATCAGATACGGTGATCCTGCCCACCTCCATGAGATCGCCAAAAACGAGGCCGTGGTGA
AGTTCGTGCTCGGCAGGATCGCTGACATCCAGATGCATCTTTTGGAACGTATCGAAAAGAATTTTGT
CATCACGGATCCGCGTCTTCCCGACAATCCGATTATCTTCGCGTCAGACTCTTTCTTACAACTGACTG
AGTATAGTAGAGAGGAGATATTGGGGCGTAACTGTAGATTTCTTCAGGGGCCAGAAACTGATCGGG
CTACCGTTCGCAAGATACGTGACGCAATAGACAACCAGACCGAGGTGACGGTGCAGCTGATTAACT
ACACAAAGTCTGGGAAGAAGTTCTGGAACCTGTTTCATTTACAACCTATGAGAGATCAAAAAGGTGAC
GTTCAATATTTCATCGGGGTTCAGTTAGATGGGACTGAGCACGTGAGAGATGCAGCAGAAAGAGAG
GGTGTAATGCTTATTAAAAAAACAGCCGAGAATATCGACGAAGCCGCTAAGCGTCACCAGAAAAAAC
AGGGCCAGAACGGCAAGAACCAGATCGACAGGTACTACGAAACTTGTATCGGAAAGGATAAGGGCA
AGAGCGTGAGCGAAAAGGTGGACGCTCTCACAAAGATCATCACCGGAATGAACTACGACCAATTCG
ACAAGAAAAGGAGCGTCATTGAGGACACCGGCAGGGAAAACGCCGAGAGGGAGAAGTTTAAAAAGA
TCATCAGCCTGTACCTCACCGTGATCTACCACATCCTCAAGAATATTGTCAATATCAACGCCAGGTAC
GTCATCGGATTCCATTGCGTCGAGCGTGATGCTCAACTGTACAAGGAGAAAGGCTACGACATCAATC
TCAAGAAACTGGAAGAGAAGGGATTCAGCTCCGTCACCAAGCTCTGCGCTGGCATTGATGAAACTG
CCCCCGATAAGAGAAAGGACGTGGAAAAGGAGATGGCTGAAAGAGCCAAGGAGAGCATTGACAGC
CTCGAGAGCGCCAACCCCAAGCTGTATGCCAATTACATCAAATACAGCGACGAGAAGAAAGCCGAG
GAGTTCACCAGGCAGATTAACAGGGAGAAGGCCAAAACCGCCCTGAACGCCTACCTGAGGAACACC
AAGTGGAATGTGATCATCAGGGAGGACCTCCTGAGAATTGACAACAAGACATGTACCCTGTTCAGAA
ACAAGGCCGTCCACCTGGAAGTGGCCAGGTATGTCCACGCCTATATCAACGACATTGCCGAGGTCA
ATTCCTACTTCCAACTGTACCATTACATCATGCAGAGAATTATCATGAATGAGAGGTACGAGAAAAGC
AGCGGAAAGGTGTCCGAGTACTTCGACGCTGTGAATGACGAGAAGAAGTACAACGATAGGCTCCTG

AAACTGCTGTGTGTGCCTTTCGGCTACTGTATCCCCAGGTTTAAGAACCTGAGCATCGAGGCCCTGT
TCGATAGGAACGAGGCCGCCAAGTTCGACAAGGAGAAAAAGAAGGTGTCCGGCAATTCC

#### Supplementary Note 2: sequence of RfxCas13d-LightR(QK634)

In black: RfxCas13d In blue: LightR

ATCGAAAAAAAAAAGTCCTTCGCCAAGGGCATGGGCGTGAAGTCCACACTCGTGTCCGGCTCCAAA
GTGTACATGACAACCTTCGCCGAAGGCAGCGACGCCAGGCTGGAAAAGATCGTGGAGGGCGACAG
CATCAGGAGCGTGAATGAGGGCGAGGCCTTCAGCGCTGAAATGGCCGATAAAAACGCCGGCTATAA
GATCGGCAACGCCAAATTCAGCCATCCTAAGGGCTACGCCGTGGTGGCTAACAACCCTCTGTATACA
GGACCCGTCCAGCAGGATATGCTCGGCCTGAAGGAAACTCTGGAAAAGAGGTACTTCGGCGAGAGC
GCTGATGGCAATGACAATATTTGTATCCAGGTGATCCATAACATCCTGGACATTGAAAAAATCCTCGC
CGAATACATTACCAACGCCGCCTACGCCGTCAACAATATCTCCGGCCTGGATAAGGACATTATTGGA
TTCGGCAAGTTCTCCACAGTGTATACCTACGACGAATTCAAAGACCCCGAGCACCATAGGGCCGCTT
TCAACAATAACGATAAGCTCATCAACGCCATCAAGGCCCAGTATGACGAGTTCGACAACTTCCTCGA
TAACCCCAGACTCGGCTATTTCGGCCAGGCCTTTTTCAGCAAGGAGGGCAGAAATTACATCATCAAT
TACGGCAACGAATGCTATGACATTCTGGCCCTCCTGAGCGGACTGAGGCACTGGGTGGTCCATAAC
AACGAAGAAGAGTCCAGGATCTCCAGGACCTGGCTCTACAACCTCGATAAGAACCTCGACAACGAAT
ACATCTCCACCCTCAACTACCTCTACGACAGGATCACCAATGAGCTGACCAACTCCTTCTCCAAGAA
CTCCGCCGCCAACGTGAACTATATTGCCGAAACTCTGGGAATCAACCCTGCCGAATTCGCCGAACAA
TATTTCAGATTCAGCATTATGAAAGAGCAGAAAAACCTCGGATTCAATATCACCAAGCTCAGGGAAGT
GATGCTGGACAGGAAGGATATGTCCGAGATCAGGAAAAATCATAAGGTGTTCGACTCCATCAGGACC
AAGGTCTACACCATGATGGACTTTGTGATTTATAGGTATTACATCGAAGAGGATGCCAAGGTGGCTG
CCGCCAATAAGTCCCTCCCCGATAATGAGAAGTCCCTGAGCGAGAAGGATATCTTTGTGATTAACCT
GAGGGGCTCCTTCAACGACGACCAGAAGGATGCCCTCTACTACGATGAAGCTAATAGAATTTGGAGA
AAGCTCGAAAATATCATGCACAACATCAAGGAATTTAGGGGAAACAAGACAAGAGAGTATAAGAAGA
AGGACGCCCCTAGACTGCCCAGAATCCTGCCCGCTGGCCGTGATGTTTCCGCCTTCAGCAAACTCA
TGTATGCCCTGACCATGTTCCTGGATGGCAAGGAGATCAACGACCTCCTGACCACCCTGATTAATAA
ATTCGATAACATCCAGAGCTTCCTGAAGGTGATGCCTCTCATCGGAGTCAACGCTAAGTTCGTGGAG
GAATACGCCTTTTTCAAAGACTCCGCCAAGATCGCCGATGAGCTGAGGCTGATCAAGTCCTTCGCTA
GAATGGGAGAACCTATTGCCGATGCCAGGAGGGCCATGTATATCGACGCCATCCGTATTTTAGGAAC
CAACCTGTCCTATGATGAGCTCAAGGCCCTCGCCGACACCTTTTCCCTGGACGAGAACGGAAACAA
GCTCAAGAAAGGCAAGCACGGCATGAGAAATTTCATTATTAATAACGTGATCAGCAATAAAAGGTTCC
ACTACCTGATCAGATACGGTGATCCTGCCCACCTCCATGAGATCGCCAAAAACGAGGCCGTGGTGA
AGTTCGTGCTCGGCAGGATCGCTGACATCCAGggaccaggtggcagcggaggtcataccttgtatgcgccgggggttatg
acatcatgggttacctcatacagatcatgaataggccgaacccacaagtggagctcggacccgtcgatacctcctgcgctctcattctgtgtgacctta
agcagaaggataccccctatcgtgtacgcctccgaggcatttctgtacatgacagggtactcgaacgccgaagtgctgggacggaactgccgcttcc
tgcaaagcccggatggaatggtgaagcctaagtcaacccgcaaatacgtggactccaacactatcaacaccatgcgcaaggccattgaccgca
atgctgaggtgcaagtggaagtggtgaacttcaagaagaatggacagcgcttcgtcaacttcctgactatgatccccgtgcgcgacgagaccggc
gaataccggtacagcatggggtttcagtgtgagacagagggcggctccggaggcagcggcggttctggaggttccggtggcggctccggaggta
gcggagggtctcacactctttacgcccctggaggatacgacattatgggatatttgattcagattatgaaccgcccaaaccctcaggtcgaactgggg
cctgtggacacgtcatgtgccctgatcctgtgcgatctgaagcaaaaggacactccgattgtctacgcctcggaagccttcttgtatatgaccggatac
agcaatgcagaggtgctcggtaggaactgcagattcctgcagtcccccgacggatggtgaaaccaaagtcgactcgcaaatatgtggactcga
acacgatcaatacaatgcggaaggccatcgaccggaacgccgaggtccaggtggaggtggtcaactttaagaagaacggccagcggttcgtga
actttctcaccatgattccggtccgggatgaaaccggagagtacagatactccatgggattccagtgcgaaaccgaagggtccggaggtcccgga
AAAAAACAGGGCCAGAACGGCAAGAACCAGATCGACAGGTACTACGAAACTTGTATCGGAAAGGAT
AAGGGCAAGAGCGTGAGCGAAAAGGTGGACGCTCTCACAAAGATCATCACCGGAATGAACTACGAC
CAATTCGACAAGAAAAGGAGCGTCATTGAGGACACCGGCAGGGAAAACGCCGAGAGGGAGAAGTTT
AAAAAGATCATCAGCCTGTACCTCACCGTGATCTACCACATCCTCAAGAATATTGTCAATATCAACGC
CAGGTACGTCATCGGATTCCATTGCGTCGAGCGTGATGCTCAACTGTACAAGGAGAAAGGCTACGA
CATCAATCTCAAGAAACTGGAAGAGAAGGGATTCAGCTCCGTCACCAAGCTCTGCGCTGGCATTGAT
GAAACTGCCCCCGATAAGAGAAAGGACGTGGAAAAGGAGATGGCTGAAAGAGCCAAGGAGAGCATT
GACAGCCTCGAGAGCGCCAACCCCAAGCTGTATGCCAATTACATCAAATACAGCGACGAGAAGAAA

GCCGAGGAGTTCACCAGGCAGATTAACAGGGAGAAGGCCAAAACCGCCCTGAACGCCTACCTGAG
GAACACCAAGTGGAATGTGATCATCAGGGAGGACCTCCTGAGAATTGACAACAAGACATGTACCCTG
TTCAGAAACAAGGCCGTCCACCTGGAAGTGGCCAGGTATGTCCACGCCTATATCAACGACATTGCC
GAGGTCAATTCCTACTTCCAACTGTACCATTACATCATGCAGAGAATTATCATGAATGAGAGGTACGA
GAAAAGCAGCGGAAAGGTGTCCGAGTACTTCGACGCTGTGAATGACGAGAAGAAGTACAACGATAG
GCTCCTGAAACTGCTGTGTGTGCCTTTCGGCTACTGTATCCCCAGGTTTAAGAACCTGAGCATCGAG
GCCCTGTTCGATAGGAACGAGGCCGCCAAGTTCGACAAGGAGAAAAAGAAGGTGTCCGGCAATTCC

#### Supplementary Note 3: sequence of RfxCas13d-UniRapR (QK634)

In black: RfxCas13d          In brown: UniRapR

ATCGAAAAAAAAAAGTCCTTCGCCAAGGGCATGGGCGTGAAGTCCACACTCGTGTCCGGCTCCAAA
GTGTACATGACAACCTTCGCCGAAGGCAGCGACGCCAGGCTGGAAAAGATCGTGGAGGGCGACAG
CATCAGGAGCGTGAATGAGGGCGAGGCCTTCAGCGCTGAAATGGCCGATAAAAACGCCGGCTATAA
GATCGGCAACGCCAAATTCAGCCATCCTAAGGGCTACGCCGTGGTGGCTAACAACCCTCTGTATACA
GGACCCGTCCAGCAGGATATGCTCGGCCTGAAGGAAACTCTGGAAAAGAGGTACTTCGGCGAGAGC
GCTGATGGCAATGACAATATTTGTATCCAGGTGATCCATAACATCCTGGACATTGAAAAAATCCTCGC
CGAATACATTACCAACGCCGCCTACGCCGTCAACAATATCTCCGGCCTGGATAAGGACATTATTGGA
TTCGGCAAGTTCTCCACAGTGTATACCTACGACGAATTCAAAGACCCCGAGCACCATAGGGCCGCTT
TCAACAATAACGATAAGCTCATCAACGCCATCAAGGCCCAGTATGACGAGTTCGACAACTTCCTCGA
TAACCCCAGACTCGGCTATTTCGGCCAGGCCTTTTTCAGCAAGGAGGGCAGAAATTACATCATCAAT
TACGGCAACGAATGCTATGACATTCTGGCCCTCCTGAGCGGACTGAGGCACTGGGTGGTCCATAAC
AACGAAGAAGAGTCCAGGATCTCCAGGACCTGGCTCTACAACCTCGATAAGAACCTCGACAACGAAT
ACATCTCCACCCTCAACTACCTCTACGACAGGATCACCAATGAGCTGACCAACTCCTTCTCCAAGAA
CTCCGCCGCCAACGTGAACTATATTGCCGAAACTCTGGGAATCAACCCTGCCGAATTCGCCGAACAA
TATTTCAGATTCAGCATTATGAAAGAGCAGAAAAACCTCGGATTCAATATCACCAAGCTCAGGGAAGT
GATGCTGGACAGGAAGGATATGTCCGAGATCAGGAAAAATCATAAGGTGTTCGACTCCATCAGGACC
AAGGTCTACACCATGATGGACTTTGTGATTTATAGGTATTACATCGAAGAGGATGCCAAGGTGGCTG
CCGCCAATAAGTCCCTCCCCGATAATGAGAAGTCCCTGAGCGAGAAGGATATCTTTGTGATTAACCT
GAGGGGCTCCTTCAACGACGACCAGAAGGATGCCCTCTACTACGATGAAGCTAATAGAATTTGGAGA
AAGCTCGAAAATATCATGCACAACATCAAGGAATTTAGGGGAAACAAGACAAGAGAGTATAAGAAGA
AGGACGCCCCTAGACTGCCCAGAATCCTGCCCGCTGGCCGTGATGTTTCCGCCTTCAGCAAACTCA
TGTATGCCCTGACCATGTTCCTGGATGGCAAGGAGATCAACGACCTCCTGACCACCCTGATTAATAA
ATTCGATAACATCCAGAGCTTCCTGAAGGTGATGCCTCTCATCGGAGTCAACGCTAAGTTCGTGGAG
GAATACGCCTTTTTCAAAGACTCCGCCAAGATCGCCGATGAGCTGAGGCTGATCAAGTCCTTCGCTA
GAATGGGAGAACCTATTGCCGATGCCAGGAGGGCCATGTATATCGACGCCATCCGTATTTTAGGAAC
CAACCTGTCCTATGATGAGCTCAAGGCCCTCGCCGACACCTTTTCCCTGGACGAGAACGGAAACAA
GCTCAAGAAAGGCAAGCACGGCATGAGAAATTTCATTATTAATAACGTGATCAGCAATAAAAGGTTCC
ACTACCTGATCAGATACGGTGATCCTGCCCACCTCCATGAGATCGCCAAAAACGAGGCCGTGGTGA
AGTTCGTGCTCGGCAGGATCGCTGACATCCAGggaccaggtacctgcgtggtgcactacaccgggatgcttgaagatggaa
agaaatttgattcctcccgggacagaaacaagccctttaagtttatgctaggcaagcaggaggtgatccgaggctgggaagaaggggttgcccag
atgagtgtgggtcagagagccaaactgactatatctccagattatgcctatggtgccactgggcacggttcgggctccggatcaggcgtcaaggacc
tcctccaagcctgggacctctattatcatgtgttccgacgaatctcaggtcctccaggacctggatcaggtctctggcatgagatgtggcatgaaggcc
tggaagaggcatctcgtttgtactttggggaaaggaacgtgaaaggcatgtttgaggtgctggagcccttgcatgctatgatggaacggggcccca
gactctgaaggaaacatcctttaatcaggcctatggtcgagatttaatggaggcccaagagtggtgcaggaagtacatgaaatcagggtcatcagg
gggctccggatcaggcatcatcccaccacatgccactctcgtcttcgatgtggagcttctaaaactggaaggtcccggaAAAAAACAGGGC
CAGAACGGCAAGAACCAGATCGACAGGTACTACGAAACTTGTATCGGAAAGGATAAGGGCAAGAGC
GTGAGCGAAAAGGTGGACGCTCTCACAAAGATCATCACCGGAATGAACTACGACCAATTCGACAAGA
AAAGGAGCGTCATTGAGGACACCGGCAGGGAAAACGCCGAGAGGGAGAAGTTTAAAAAGATCATCA
GCCTGTACCTCACCGTGATCTACCACATCCTCAAGAATATTGTCAATATCAACGCCAGGTACGTCATC
GGATTCCATTGCGTCGAGCGTGATGCTCAACTGTACAAGGAGAAAGGCTACGACATCAATCTCAAGA
AACTGGAAGAGAAGGGATTCAGCTCCGTCACCAAGCTCTGCGCTGGCATTGATGAAACTGCCCCCG
ATAAGAGAAAGGACGTGGAAAAGGAGATGGCTGAAAGAGCCAAGGAGAGCATTGACAGCCTCGAGA

GCGCCAACCCCAAGCTGTATGCCAATTACATCAAATACAGCGACGAGAAGAAAGCCGAGGAGTTCA
CCAGGCAGATTAACAGGGAGAAGGCCAAAACCGCCCTGAACGCCTACCTGAGGAACACCAAGTGGA
ATGTGATCATCAGGGAGGACCTCCTGAGAATTGACAACAAGACATGTACCCTGTTCAGAAACAAGGC
CGTCCACCTGGAAGTGGCCAGGTATGTCCACGCCTATATCAACGACATTGCCGAGGTCAATTCCTAC
TTCCAACTGTACCATTACATCATGCAGAGAATTATCATGAATGAGAGGTACGAGAAAAGCAGCGGAA
AGGTGTCCGAGTACTTCGACGCTGTGAATGACGAGAAGAAGTACAACGATAGGCTCCTGAAACTGCT
GTGTGTGCCTTTCGGCTACTGTATCCCCAGGTTTAAGAACCTGAGCATCGAGGCCCTGTTCGATAGG
AACGAGGCCGCCAAGTTCGACAAGGAGAAAAAGAAGGTGTCCGGCAATTCC
